# FkpA Enhances Membrane Protein Folding using an Extensive Interaction Surface

**DOI:** 10.1101/2022.11.01.514694

**Authors:** Taylor A. Devlin, Dagan C. Marx, Michaela A. Roskopf, Quenton R. Bubb, Ashlee M. Plummer, Karen G. Fleming

## Abstract

Outer membrane protein (OMP) biogenesis in gram-negative bacteria is managed by a network of periplasmic chaperones that includes SurA, Skp, and FkpA. These chaperones bind unfolded OMPs (uOMPs) in dynamic conformational ensembles to suppress uOMP aggregation, facilitate diffusion across the periplasm, and enhance OMP folding. FkpA primarily responds to heat-shock stress, but its mechanism is comparatively understudied. To determine FkpA chaperone function, we monitored the folding of a cognate client uOmpA_171_ and found that FkpA increases the folded uOmpA_171_population but also slows the folding rate, dual functions distinct from the other periplasmic chaperones. The results indicate that FkpA behaves as a chaperone and not as a folding catalyst to directly influence the uOmpA_171_folding trajectory. We determine the binding affinity between FkpA and uOmpA_171_ by globally fitting sedimentation velocity titrations and found it to be intermediate between the known affinities of Skp and SurA for uOMP clients. Notably, complex formation steeply depends on the urea concentration, suggestive of an extensive binding interface. Initial characterizations of the complex using photo-crosslinking indicates that the binding interface spans the inner surfaces of the entire FkpA molecule. In contrast to prior findings, folding and binding experiments performed using subdomain constructs of FkpA demonstrate that the full-length chaperone is required for full activity. Together these results support that FkpA has a distinct and direct effect on uOMP folding and that it achieves this by utilizing an extensive chaperone-client interface.

**Significance:** The periplasmic chaperone network is required for the survival and virulence of gram-negative bacteria. Here we find that the chaperone FkpA enhances outer membrane protein folding and tightly binds its clients with an extensive interaction interface. This modified holdase function of FkpA distinguishes it from other periplasmic chaperones and complements their functions to ensure robust outer membrane biogenesis.

## Introduction

The outer membrane of gram-negative bacteria serves as the first line of defense against extracellular threats, and outer membrane biogenesis and stability are essential for cell viability.^1^ The outer membrane is composed of lipids and outer membrane proteins (OMPs) that facilitate the flux of small molecules, maintain membrane integrity, and promote virulence in pathogenic strains. To fold into the outer membrane, an OMP must first traffic across multiple cellular compartments in an unfolded but folding-competent conformation.^2^ Unfolded OMPs (uOMPs) are translated in the cytoplasm and post-translationally translocated into the aqueous periplasm through the Sec YEG translocon.^3^ Because uOMPs contain highly hydrophobic regions and otherwise readily aggregate in water^4,5^ they are bound and released by a network of periplasmic chaperones to prevent off-pathway aggregation before finally folding into the outer membrane catalyzed by the β-barrel assembly machine (BAM).^2,6^ Importantly, all of these processes must occur in the absence of ATP^7^ and be primed to respond to changing environmental conditions that could occur as a consequence of the leaky outer membrane. How do periplasmic chaperones suppress uOMP aggregation and promote native folding in the absence of an external energy source and in the face of external stress? Understanding the general principles governing these ATP-independent chaperone-client interactions contributes to the growing comprehension of bacterial protein homeostasis in virulent strains and informs the development of antibiotics targeting the cell envelope.

Major players in the OMP biogenesis chaperone network include SurA, Skp, and FkpA with additional contributions from Spy and the protease chaperone DegP (Figure 1).^8,9^ While much of the recent work elucidating the mechanisms of these ATP-independent chaperones focuses on the chaperones SurA and Skp,^10,11,20,21,12–19^ thermodynamic and mechanistic information about the function of FkpA in OMP biogenesis remains sparse.^22,23^ Structurally, FkpA forms homodimers and comprises two structural domains: an N-terminal, α-helical domain that mediates dimerization and a C-terminal domain homologous to FK506-binding proteins and capable of catalyzing peptidyl-prolyl isomerization reactions.^24–27^ The structural domains are connected by a long and flexible α-helix, allowing the two C-terminal domains to move independently of each other.^27–29^ There are unstructured tails at both the N- and C-termini, but these intrinsically disordered regions have not been ascribed any functional importance.^27,30^

**Figure 1.**
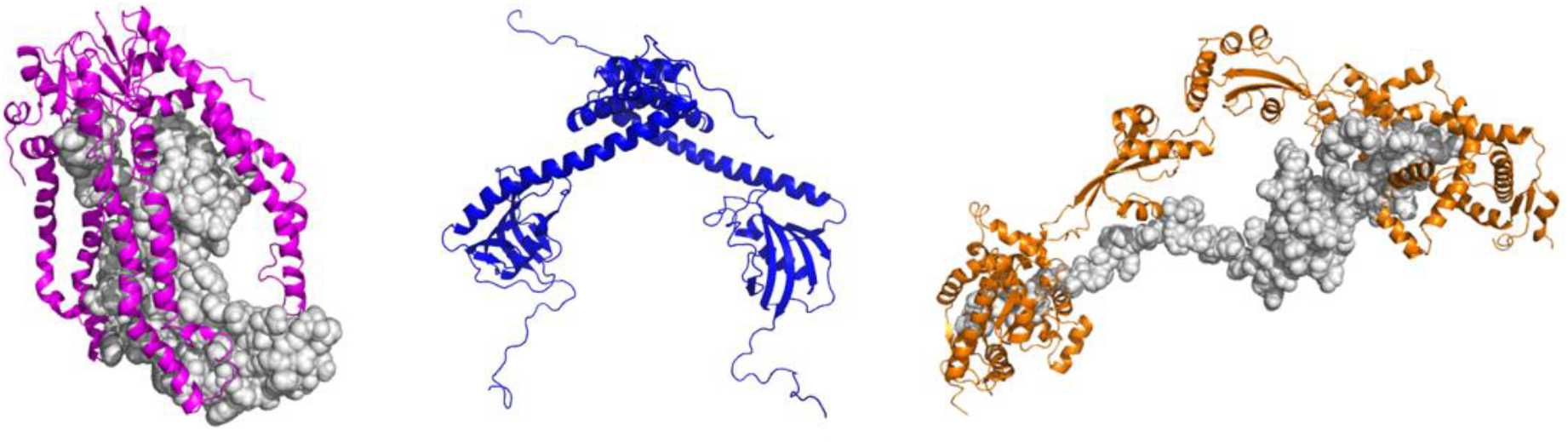
Three periplasmic chaperones involved in OMP biogenesis. Well studied periplasmic chaperones Skp (left, pink) and SurA (right, orange) use distinct mechanisms to chaperone unfolded outer membrane proteins in the periplasm, and models of each complex have been published. By contrast, limited information about the mechanism and chaperone-client complex is available for FkpA (middle, blue). The dimeric structure of FkpA was created from PDB 1Q6U^27^ with the N- and C-terminal tails added in PyMol.^72^ The model of Skp binding uOmpA_171_ is modified from Zaccai *et al*.^66^ The model of multiple SurA molecules binding an extended uOmpA_171_ is modified from Marx *et al*.^10^

FkpA manifests its chaperone function by increasing the solubility and decreasing aggregation of a variety of unfolded or misfolded proteins (including non-OMPs) *in vitro*.^24–28,31^ Additionally, FkpA improves yields of recombinantly expressed proteins when co-expressed in the same system.^25,32–36^ In the context of OMP biogenesis, FkpA rescues *ΔsurA Δskp* lethality at elevated temperatures^22^ and is upregulated by the *E. coli* sigma E (σ^E^) stress response,^37,38^ implying this chaperone contributes as an important mitigator of increased uOMP aggregation risk in the periplasm under stress conditions. Due to the redundant and functionally overlapping nature of the OMP biogenesis chaperone network, FkpA is not essential to cell survival, but cumulative data suggest that it is a potent stress-response chaperone with a wide range of possible OMP and non-OMP clients.

Despite evidence for FkpA exhibiting chaperone function on model unfolded or misfolded proteins both *in vivo* and *in vitro*, limited information is available about FkpA binding to physiologically relevant OMPs^22,23^. It is also not known how FkpA influences the folding trajectory of uOMPs. Here we address these questions using the transmembrane β-barrel of outer membrane protein A (OmpA_171_) to study both the chaperone function and binding affinity of FkpA. The full-length and dimeric FkpA exhibits chaperone function in an OmpA_171_ folding assay by increasing the total folded population of OmpA_171_ but at the same time reduces the rate of OmpA_171_ folding. This chaperone activity is distinct from the effects of periplasmic chaperones SurA or Skp. FkpA tightly binds uOmpA_171_ with low nM affinity in the absence of denaturant. We observe a steep urea-dependence of the binding constant indicating that a large surface area is buried upon client binding. We find the FkpA binding affinity for uOMPs to be more favorable than that of SurA but less favorable than that of Skp based on direct competition for uOmpA_171_ binding.

Because FkpA has two distinct structural domains, it is tempting to hypothesize that each domain makes distinct contributions to client-binding and chaperone function. Previous work has sought to elucidate these domain contributions, but the prior data do not come to a consensus view. Some data indicate that the N-terminal dimerization domain is not only required but sufficient for chaperone function,^26,27^ but other results localize the client-binding and chaperone function to the C-terminal peptidyl-prolyl isomerase (PPIase) domains.^24,28^ To reconcile these observations, we tested the chaperone function and binding affinity of the individual domains, the N-terminal dimerization domain (N-FkpA) and the C-terminal PPIase domain (C-FkpA), and compared these results to those of full-length FkpA. We find that individual domain alone neither exhibits chaperone function nor accounts for the full binding affinity. Supporting this conclusion, site specific photo-crosslinking between FkpA and uOmpA_171_ indicates that the interaction interface spans both domains. We hypothesize that dimerization mediated by the N-domain brings the two PPIase domains into close proximity to create an extensive binding surface with much higher affinity for unfolded or misfolded clients.

## Results

### FkpA Increases the Folded Fraction but Slows the Folding Rate of OmpA_171_

We investigated the function of FkpA (FkpA refers to the full-length, dimeric form of the protein unless otherwise stated) by observing its effect on the folding the β-barrel domain of outer membrane protein A (OmpA_171_). This eight-stranded β-barrel is a useful client OMP to study FkpA chaperone function because it is abundant in the outer membrane of *E. coli*, it contains no *cis*-prolines (precluding confounding effects from the known PPIase activity of FkpA), and its folding pathway has been thoroughly investigated *in vitro*.^39,40^ We take advantage of the heat-modifiability of folded OMPs in mixed SDS-phospholipid micelles to separate folded and unfolded populations into distinct bands on an SDS-PAGE gel that can be quantified by densitometry.^41–46^ As has been demonstrated previously in the literature, the folded and unfolded fractions do not add to unity due to the presence of an elusive state that migrates anomalously on a polyacrylamide gel.^40^ Therefore, a fraction elusive was also quantified. The exact nature of this elusive state is unknown, but OmpA_171_ has been shown to form unfolded oligomers in the presence of lipids or detergents,^47^ and this state represents off-pathway, lipid-induced aggregates in this folding assay.^40^ Full details are provided in the Supplemental Materials and Methods.

In the absence of chaperone, roughly 50% of the total OmpA_171_ folds into 1,2-diundecanoyl-_sn_-glycero-3-phosphocholine (diC_11_PC) LUVs after one hour (Figure 2AB, SI Figure S1) in excellent agreement with previous studies of OmpA_171_ folding into the same lipid environment.^40,46^ The addition of FkpA to the folding reaction significantly increases the folded fraction of OmpA_171_ to ~70% (Figure 2AB, SI Figure S1), demonstrating that this periplasmic chaperone has a direct beneficial effect on OmpA_171_ folding. Notably, this improved folding in the presence of FkpA is unique; other periplasmic chaperones do not have this effect. SurA, presumably the most utilized periplasmic chaperone in OMP biogenesis,^48,49^ does not impact OmpA_171_ folding in this assay (Figure 2A-C) while Skp maintains the protein almost entirely in the unfolded state over the course of an hour (Figure 2ABD) in agreement with previously published results.^13,21^ Together these observations and comparisons to other players in the OMP biogenesis pathway differentiates FkpA and indicates that it uses a distinct mechanism to improve OMP folding.

**Figure 2.**
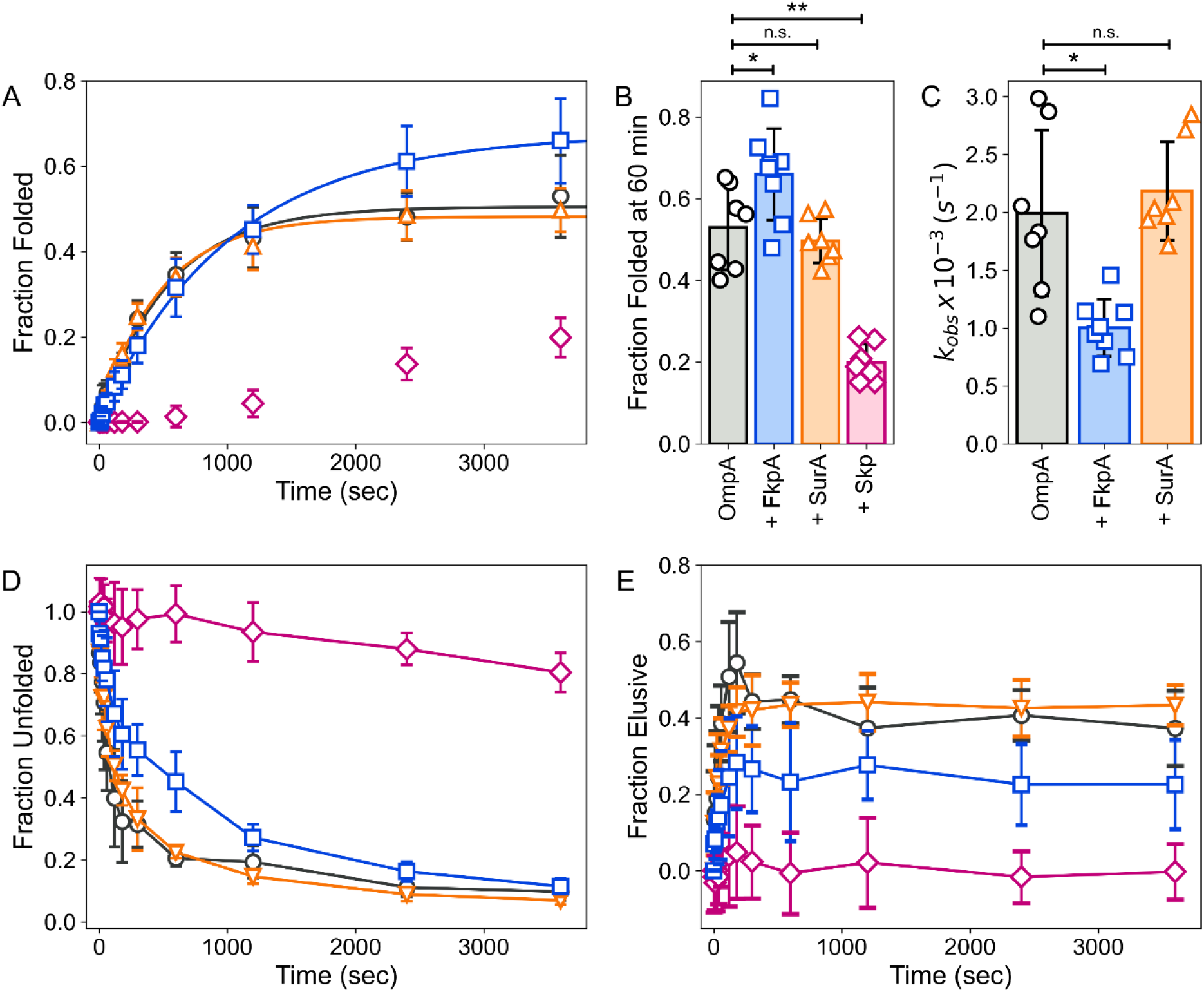
FkpA increases the folded population of OmpA_171_ at 60 min but slows the observed rate of OmpA_171_ folding. A) Fraction of OmpA_171_ folded into diC_11_PC LUVS as a function of time (0-3600 s) in the presence of FkpA (blue squares, n=8), SurA (orange triangles, n=7), or Skp (pink diamonds, n=7) or without any chaperone (black circles, n=7). Error bars represent ±1 standard deviation. Lines are single exponential fits to the average data. Folding in the presence of Skp is not fit to an exponential function due to the long lag period. B) Comparison of the fraction of OmpA_171_ folded at 60 min. C) Comparison of the observed rate constants when individual datasets are fit to single exponential functions. (* p < 0.05, ** p<0.01, independent t-test with unequal variances) D) Unfolded fraction of OmpA_171_ as a function of time. E) Elusive fraction as a function of time. Error bars represent ±1 standard deviation.

To gain insight into the effect of FkpA on the kinetics of OmpA_171_ folding, the folding profiles were fit to single exponential functions to obtain observed first-order rate constants (k_obs_). For OmpA_171_ alone, k_obs_ = 2.0 (± 0.7) x 10^−3^ s^−1^(Figure 2C), which is in good agreement with the previously observed slow component of OmpA_171_ folding.^46^ Upon adding FkpA to the folding reactions, the observed OmpA_171_ folding rate decreases by a factor of two (k_obs_ = 1.0 (± 0.2) x 10^−3^ s^−1^, Figure 2C). The decrease in OmpA_171_ folding rate in the presence of FkpA indicates that FkpA behaves as a uOMP chaperone and not as a folding catalyst, since folding catalysts (like BamA for example^50^) increase the rate of uOMP folding. The lack of *cis*-prolines in the primary sequence of OmpA_171_ ensures that the known PPIase activity of FkpA plays no role in improving the folding of OmpA_171_. In fact, *cis*-prolines are generally uncommon among cell envelope proteins (SI Table S1), suggesting that the chaperone function of FkpA supersedes its PPIase activity in the periplasm. Thus, both the increase in the fraction of OmpA_171_ folded and the decrease in folding rate are attributed to the chaperone function of FkpA.

Beyond changes to the folded population of OmpA_171_, following the unfolded and elusive populations as a function of time provides some insight as well. FkpA maintains a larger fraction of OmpA_171_ in an unfolded state at intermediate timepoints (5-10 minutes) and decreases the final elusive population at 1 hour (Figure 2DE). Together the folding assay results suggest FkpA balances client-binding to suppress aggregation and client-release to allow folding. This keeps a larger proportion of OmpA_171_ unfolded but folding-competent at early-to-intermediate timepoints and helps prevent the formation of non-productive, off-pathway folding aggregates characteristic of the elusive state.

### FkpA Binds uOmpA_171_ with High Affinity

The chaperone function of FkpA has been demonstrated using a plethora of unfolded or misfolded model clients,^24–28,31^ but the FkpA binding affinity for known, physiologically-relevant uOMP clients is limited. To measure the affinity of FkpA for uOmpA_171_, we titrated 5 μM uOmpA_171_ with increasing FkpA concentrations between 0.01-100 μM and followed the interacting system in solution using a series of sedimentation velocity analytical ultracentrifugation experiments (SV-AUC). uOmpA_171_ is monomeric and monodisperse at a concentration of 5 μM in 1 M urea^5,51^ while FkpA is dimeric across the experimental concentrations and conditions used in this study (see *Oligomeric States of FkpA Constructs* in the Supplemental Results, SI Figure S2, SI Figure S3). Analysis of these titrations revealed the formation of an FkpA_2_*uOmpA_171_ chaperone-client complex. Complex formation is illustrated as both a concentration-dependent shift in the reaction boundary (Figure 3A) and as the change in average s_20,w_ (<s_20,w_>) as a function of FkpA concentration (SI Figure S4). The overlaid g(s*) distributions shows the depletion of the unbound uOmpA_171_ population (s_20,w_ = 1.59 Svedbergs) and formation of the chaperone-client complex (s_20,w_ = 3.80 Svedbergs) as uOmpA_171_ is titrated with FkpA. The reaction boundary and <s_20,w_> shifts back toward the distribution of unbound FkpA_2_ (s_20,w_ = 3.05 Svedbergs) due to an increasing population of free FkpA_2_ at high FkpA concentrations.

**Figure 3.**
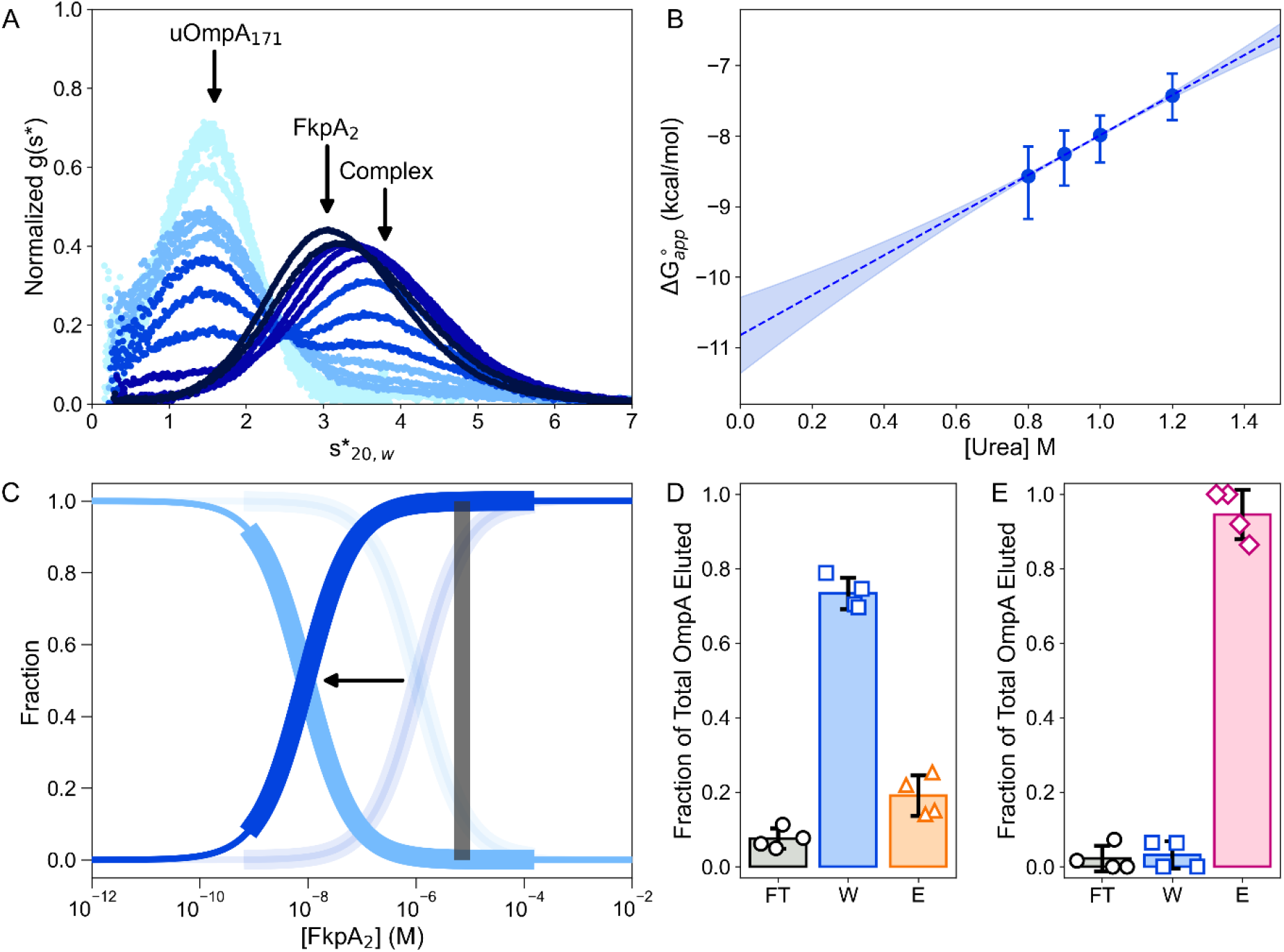
Binding of FkpA to uOmpA_171_. A) FkpA titrating 5 μM uOmpA_171_ is shown as the shift in the reaction boundary. The change in g(s*) distributions shows the depletion of the unbound uOmpA_171_ population (s_20,w_ = 1.59 (1.57-1.61) Svedbergs) and formation of the chaperone-client complex (s_20,w_ = 3.8 (3.68-3.92) Svedbergs) as uOmpA_171_ is titrated with FkpA. The reaction boundary shifts back toward the distribution of unbound FkpA_2_ due to an increasing population of free fl-FkpA_2_ at high fl-FkpA concentrations. B) Plot of the binding ΔG°_app_ as a function of urea concentration. A linear fit to the data (y = 2.8x-10.8) allows extrapolation back to 0 M urea where ΔG°_app_ = −10.8 (−10.3 - −11.3) kcal/mol and K_d_ = 8 (4-20) nM. Error bars are 95% confidence intervals on the fit calculated using the F-stat module in SEDANAL. C) The transparent fraction bound and fraction free curves represent the 1 M urea condition. The inflection point shifts leftward by two orders of magnitude in 0 M urea. Fraction of FkpA bound as a function of FkpA_2_ concentration, which assumes an obligate dimer at all concentrations, is in blue. Fraction free is light blue. Periplasmic FkpA concentrations are shown in the gray box. In the absence of all other factors, FkpA would be almost entirely bound at its periplasmic concentrations. (D) and (E) FkpA outcompetes SurA for uOmpA_171_ binding but cannot remove uOmpA_171_ bound to Skp. SurA:uOmpA_171_ or Skp:uOmpA_171_complexes were pre-equilibrated on Ni NTA resin before addition of untagged FkpA. D) Since FkpA has a greater affinity for uOmpA_171_ than SurA, the majority of the uOmpA_171_ washed off the resin with FkpA. E) uOmpA_171_ remained bound to Skp in the presence of FkpA, indicating that Skp has the stronger affinity of the two. Bars are the average of four pulldown experiments, and error bars are ± 1 standard deviation.

To determine the s_20,w_ of the FkpA_2_*uOmpA_171_ complex and the disassociation constant (K_d_) for the binding equilibrium, the full titrations were globally fit in SEDANAL (SI Figure S5) using the model:

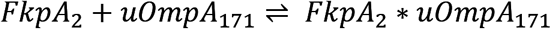

The model assumes a dimeric species of FkpA, which we and others have shown to be the dominant form of FkpA above 1 μM in concentration (see *Oligomeric States of FkpA Constructs* in the Supplemental Results),^26,31^ and also includes a small fraction of irreversible aggregate that accounts for < 1% of the total protein concentration. Sedimentation coefficients for FkpA_2_ and uOmpA_171_ were held constant at values obtained from fitting each component alone (Table 1). Globally fitting the titration in 1 M urea results in a K_d_=1.1 (0.6-1.8) μM and a complex s_20,w_ = 3.80 (3.68-3.92) Svedbergs. The 95% confidence intervals are reported in parentheses.

**Table 1.**
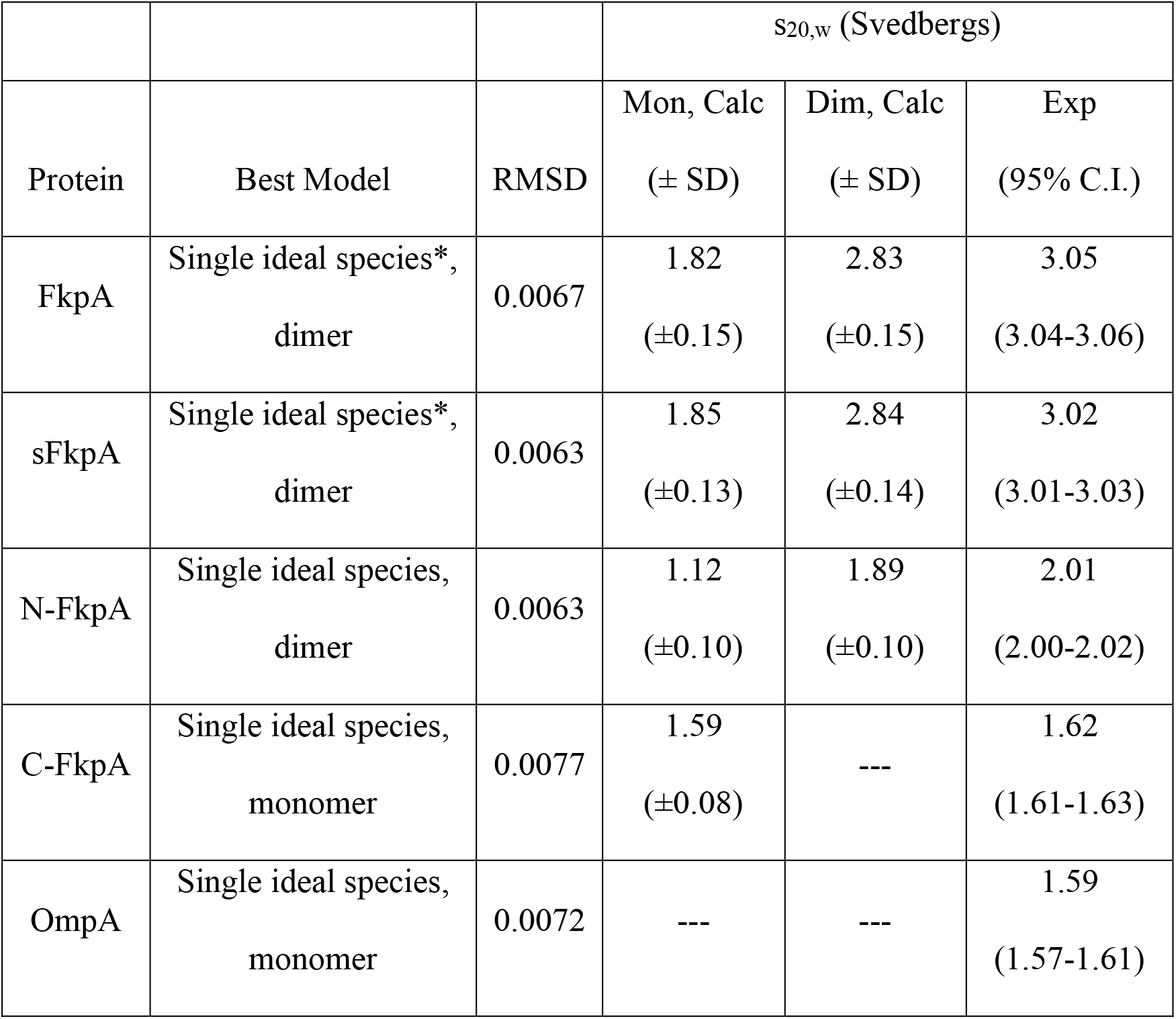
Oligomeric States and Sedimentation Coefficients of All FkpA Constructs

Just as chemical denaturants destabilize the native, folded structures of proteins, we speculate that even the low concentrations of urea in our binding titrations modulate the chaperone-client interaction. To investigate the suspected urea dependence of the binding affinity between FkpA and uOmpA_171_, we repeated titrations as a function of the urea concentration. The range of urea concentrations accessible to this type of experiment is narrow as uOmpA_171_ solubility decreases with decreasing urea concentrations,^51^ and FkpA begins to unfold above 1.5 M urea (SI Figure S6). Considering these two boundaries, we repeated the titrations in 0.8 M, 0.9 M, and 1.2 M Urea. These additional SV-AUC titrations reveal that the binding interaction between FkpA and uOmpA_171_ depends strongly on the urea concentration, but the sedimentation coefficient of the complex is relatively insensitive to the urea concentration (Table 2, Figure 3C, SI Figure S7). Analogous to the linear extrapolation method for determining protein stability from denaturation experiments, converting K_d_(urea) to ΔG_app_(urea) allows for linear extrapolation of the binding free energy change to 0 M urea (Figure 3B). The apparent K_d_ moves to a more favorable value equal to 8 (4-20) nM when extrapolated back to water. A binding constant in the low nM range qualitatively matches two previously published binding constants.^22,23^ Figure 3C illustrates the dramatic effect that even low urea concentrations have on the FkpA-uOmpA_171_ population distribution by comparing fractional binding curves in 0 and 1 M urea.

**Table 2.** Best fit parameters from global fits of FkpA SV-AUC uOmpA_171_ binding titrations

| Protein | [Urea] | Best Model | RMSD | Complex $s_{20,w}$<br>(Svedbergs)<br>(95% C.I.) | $K_d$ (μM)<br>(95% C.I.) |
| --- | --- | --- | --- | --- | --- |
| FkpA | 0.8 M | $\text{FkpA}_2 + \text{uOmpA}_{171} \leftrightarrow \text{FkpA}_2 * \text{uOmpA}_{171}^\dagger$ | 0.0078 | 3.79<br>(3.69-3.90) | 0.4<br>(0.1-0.8) |
| | 0.9 M | $\text{FkpA}_2 + \text{uOmpA}_{171} \leftrightarrow \text{FkpA}_2 * \text{uOmpA}_{171}^\dagger$ | 0.0081 | 3.73<br>(3.63-3.81) | 0.7<br>(0.3-1.2) |
| | 1.0 M | $\text{FkpA}_2 + \text{uOmpA}_{171} \leftrightarrow \text{FkpA}_2 * \text{uOmpA}_{171}^\dagger$ | 0.0071 | 3.80<br>(3.68-3.92) | 1.1<br>(0.6-1.8) |
| | 1.2 M | $\text{FkpA}_2 + \text{uOmpA}_{171} \leftrightarrow \text{FkpA}_2 * \text{uOmpA}_{171}^\dagger$ | 0.0072 | 3.86<br>(3.64-4.04) | 2.9<br>(1.6-4.9) |
| sFkpA | 1.0 M | $\text{sFkpA}_2 + \text{uOmpA}_{171} \leftrightarrow \text{sFkpA}_2 * \text{uOmpA}_{171}^\dagger$ | 0.0081 | 4.23<br>(4.16-4.30) | 3.0<br>(2.6-3.6) |
| N-FkpA | 1.0 M | $\text{N-FkpA}_2 + \text{uOmpA}_{171} \leftrightarrow \text{N-FkpA}_2 * \text{uOmpA}_{171}$ | 0.0058 | 2.73<br>(2.52-3.03) | 44<br>(28-68) |
| C-FkpA | 1.0 M | Two ideal species | 0.0072 | --- | --- |

<sup>†</sup> Includes irreversible aggregate at < 1% of the total protein concentration

The steep urea dependence of the FkpA binding affinity (m-value = 2.8 kcal*L/mol^2^) suggests that an extensive binding interface is buried upon client-binding. Using the empirical correlation between urea *m*-value and the change in accessible surface area (ΔASA) published by Myers, Pace, and Scholtz,^52^ we calculate that FkpA binding to uOmpA_171_ buries approximately 22,000 Å^2^ of ASA. The FkpA dimer alone exposes 40,000 Å^2^ of ASA while a model of uOmpA_17110_ exposes 20, 000 Å^2^ of ASA (values calculated using the FreeSASA Python module^53^). Of the sum total 60,000 Å^2^ accessible to the two unbound proteins, about 1/3 is buried in buried in the binding interface. This significant ΔASA suggests that the chaperone-client interaction involves an extensive binding interface on FkpA capable of shielding an equivalently large surface on the client from solvent.

### The Periplasmic Chaperone Binding Hierarchy

FkpA exhibits very favorable client-binding with a disassociation constant in the low nM range when extrapolated to 0 M urea. This binding affinity is stronger than affinities reported for SurA^13,54,55^ but weaker than or similar to affinities reported for Skp.^18,55,56^ To investigate the interactions between periplasmic chaperones in competition for a uOMP client, we performed competition pulldown assays based on the protocol described by Thoma *et al*.^21^ His-tagged SurA or Skp was premixed with uOmpA_171_ and added to a small batch of Ni NTA resin. Then the original chaperone-uOmpA_171_ complex was washed with untagged FkpA before eluting with high imidazole buffer. If FkpA competes the uOmpA_171_ away from the original chaperone, uOmpA_171_ appears mostly in the wash fraction. If the original chaperone binds uOmpA_171_ tighter than FkpA does, uOmpA_171_ predominantly appears in the elution fractions. FkpA does outcompete SurA for uOmpA_171_ binding, but cannot pull the uOmpA_171_ away from Skp (Figure 3DE, SI Figure S8). These results confirm our direct binding data that FkpA binds uOMPs more favorably than SurA, but Skp is the tightest binding of the three chaperones.

### OmpA_171_ Crosslinks across the Extensive Inner Surface of Dimeric FkpA

Direct structural characterization of the FkpA-uOmpA_171_ complex is extremely challenging due to the heterogeneous and dynamic nature of the unfolded OMP. We therefore approached this quest by incorporating the zero-length photo-crosslinker *para*-azido phenylalanine (pAzF) at 24 surface-exposed, non-conserved positions in FkpA using amber suppression.^57^ As had been described previously,^10^ uOmpA_171_ and an FkpA pAzF variant were mixed, exposed to ultraviolet (UV) light, and analyzed by SDS-PAGE. Crosslinking efficiency at each position was calculated using the loss of uOmpA_171_ band intensity after 5 minutes of UV exposure. An example of an SDS-PAGE readout from this crosslinking assay using the FkpA pAzF variant Y225pAzF is shown in Figure 4A. Y225 is located on the C-domain of FkpA adjacent to the PPIase catalytic site.^27^ FkpA Y225pAzF is a high efficiency crosslinking variant that crosslinks 83.2 ± 1.5 % of the original uOmpA_171_ population, shown in Figure 4A as the complete loss of the uOmpA_171_ band intensity (marked by ●) after exposure to UV light. Several higher molecular weight bands appear after UV exposure, including bands corresponding to the FkpA dimer (57 kDa, ▲▲), a single FkpA monomer crosslinked to uOmpA_171_ (37 kDa, ▲●), and an FkpA dimer crosslinked to uOmpA_171_ (66 kDa, ▲▲●). Larger crosslinked species can also form, suggesting that multiple FkpA dimers may chaperone a single uOMP client. By quantifying the crosslinking efficiencies of all 24 FkpA pAzF variants (Figure 4B, SI Figure S9) and mapping these crosslinking efficiencies on the crystal structure of FkpA (PDB 1Q6U, Figure 5), it becomes clear that the chaperone uses an extensive binding interface that spans the inner surfaces of both the N- and C-domains. This large binding interface is consistent with the steep urea dependence of the disassociation constant and suggests that several local interactions contribute to the overall tight binding affinity.

**Figure 4.**
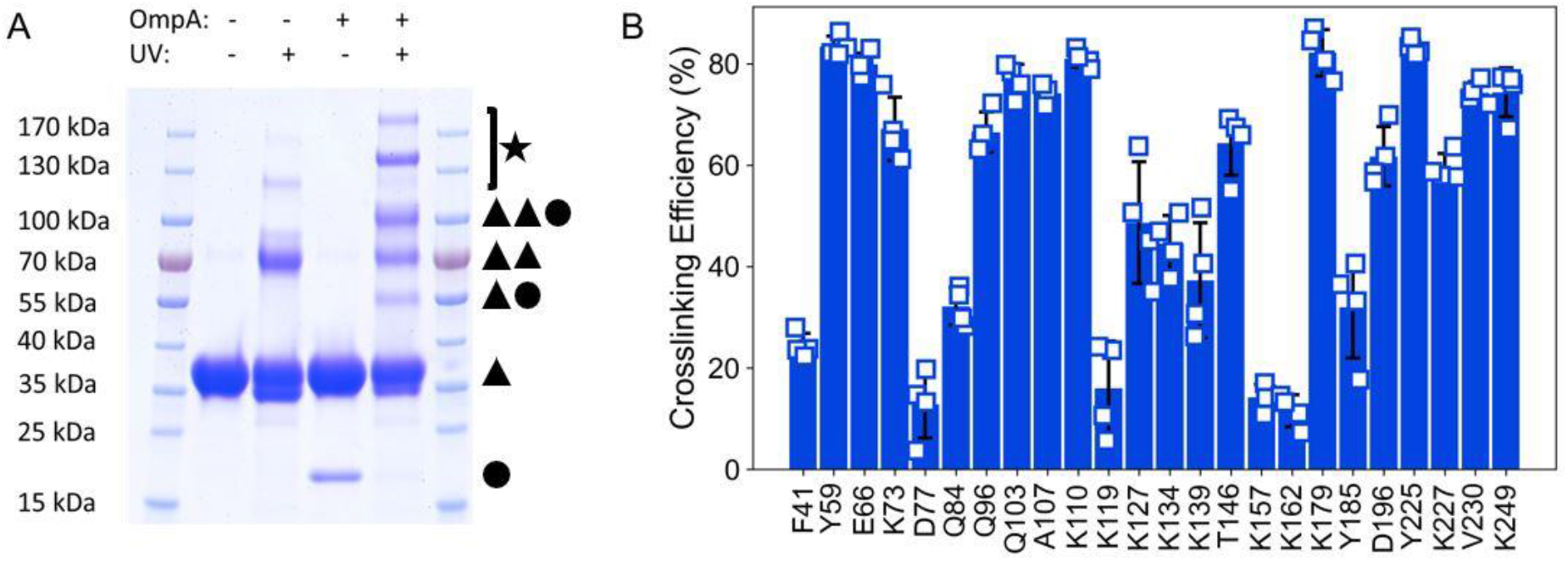
The binding interface of FkpA spans both the N- and C-domains. A) SDS-PAGE readout after photo-crosslinking FkpA variant Y225pAzF with uOmpA_171_. Y225pAzF monomers are labeled with a black triangle (▲) and uOmpA_171_ is labeled as a black circle (●). Covalently cross-linked Y225pAzF dimers formed after UV exposure are labeled using two black triangles (▲▲). Chaperone-client complexes that become photo-crosslinked include Y225pAzF:uOmpA_171_in 1:1 (▲●) and 2:1 (▲▲●) ratios (*i.e.* a monomer of FkpA and a dimer of FkpA photo-crosslinked to the OMP). Higher order complexes of Y225pAzF alone and Y225pAzF photo-crosslinked to uOmpA_171_ (black star) suggest that multiple FkpA dimers can interact with each other and with an unfolded OMP. B) Crosslinking efficiencies calculated using the loss of uOmpA_171_ band intensity for each of the 24 FkpA pAzF variants (n=4, errors bars = ±1 standard deviation).

**Figure 5.**
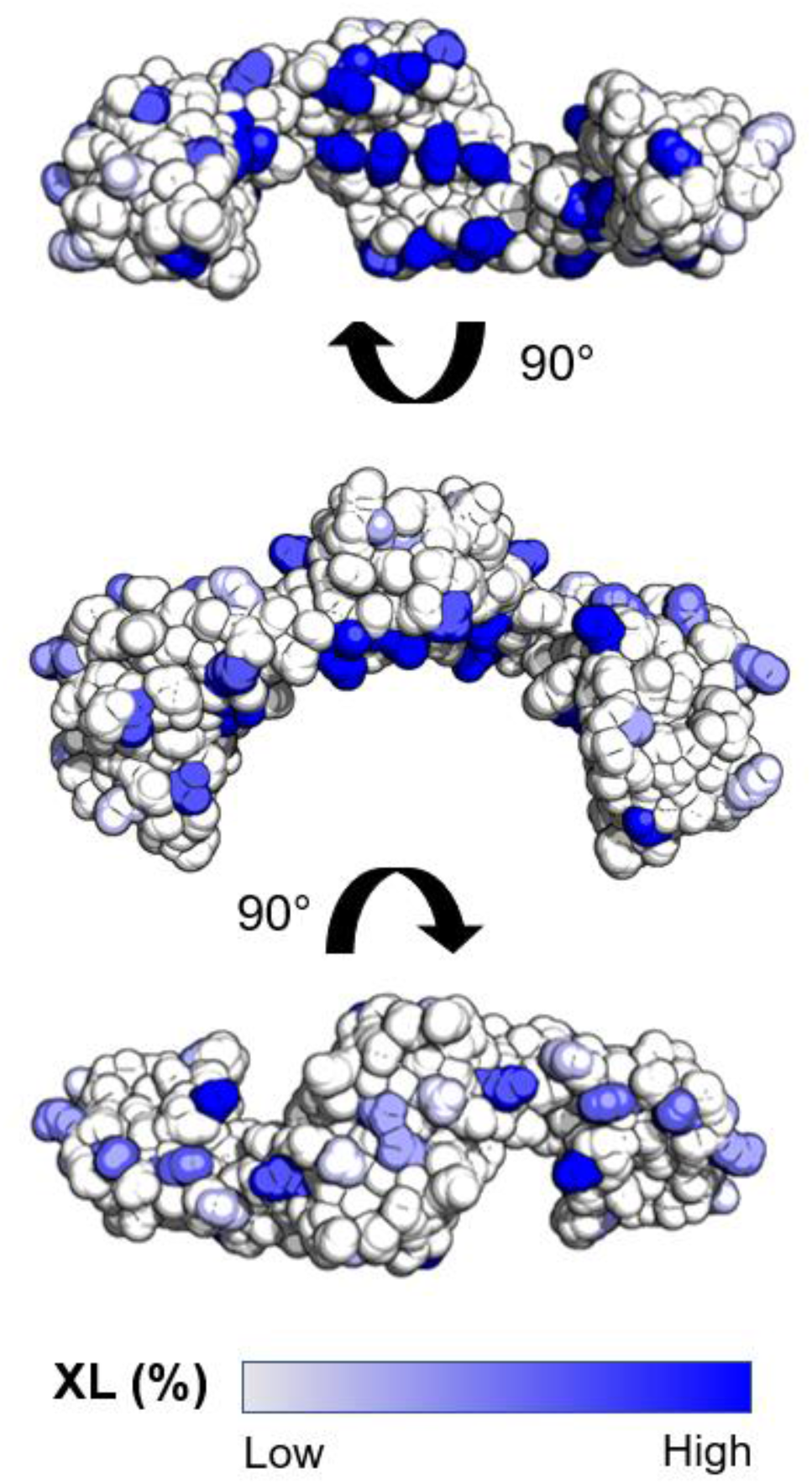
Crosslinking efficiencies shown on the surface of FkpA (PDB 1Q6U). The highest efficiency crosslinking positions are located on the inner surface between the arms of FkpA. High efficiency crosslinking positions are found on both the N- and C-domains.

### Both Structural Domains are Necessary for Tight Binding and Chaperone Function

Previous investigations sought to identify which regions of FkpA contribute to its chaperone function, but conclusions are inconsistent.^24,26–28^ Because FkpA comprises two distinct structural domains, an N-terminal dimerization domain and a C-terminal peptidyl prolyl isomerase domain, it is tempting to hypothesize that one domain contributes more to binding affinity and chaperone function than the other. However, our photo-crosslinking results suggest that surfaces on both domains contribute to the full binding interface. To further investigate the contributions of the individual domains, we created constructs of the N-terminal domain (N-FkpA) and C-terminal domain (C-FkpA). Both individual domains fold independently, and N-FkpA is dimeric while C-FkpA is monomeric (see *Oligomeric States of FkpA Constructs* in the Supplemental Results, SI Figure S2). When added to OmpA_171_ folding assays, neither domain construct had an effect on the amplitude or rate of OmpA_171_ folding, indicating that neither domain alone is sufficient to recapitulate FkpA chaperone function but that both domains are necessary (SI Figure S10). The OmpA_171_ folding experiments were also performed in the presence of sFkpA, which has the intrinsically disordered tails at the N- and C-termini of FkpA removed. Previous studies suggest that these IDRs have no functional importance *in vitro*^27,28^ or *in vivo*^22^, and while results from OmpA_171_ folding in the presence of sFkpA are not significantly different from OmpA_171_ folding alone (SI Figure S10), sFkpA does follow the same trends as full-length FkpA to slightly increase OmpA_171_ folding efficiency and slow the OmpA_171_ folding rate. Therefore, we cannot draw any conclusions about the functional importance of the IDRs from these results.

The OmpA_171_ folding assays revealed that neither FkpA domain alone demonstrates the full chaperone activity, so we evaluated their contribution to uOmpA_171_ binding using SV-AUC titrations as well. Both sFkpA and N-FkpA appear to bind uOmpA_171_ based on the comparison of calculated and experimental g(s*) distributions (Figure 6A-C). The fit binding constants and sedimentation coefficients are reported in Table 2. sFkpA binds OmpA_171_ with a K_d_ = 3.0 (2.6-3.6) μM in 1 M urea, indicating that the intrinsically disordered tails have no effect on binding affinity when compared to full-length FkpA under the same experimental conditions. N-FkpA weakly binds with a K_d_ of 44 (28-68) μM. However, calculated and experimental g(s*) distributions overlay for C-FkpA binding to uOmpA_171_, indicating no interaction (Figure 6D). Indeed, this titration was best fit by a two ideal species model, and therefore, any affinity between C-FkpA and uOmpA_171_, if it exists, is too weak to be measured under these experimental conditions. Together these results indicate that while N-FkpA has some affinity for uOmpA_171_, both domains are required to achieve the tightest binding.

**Figure 6.**
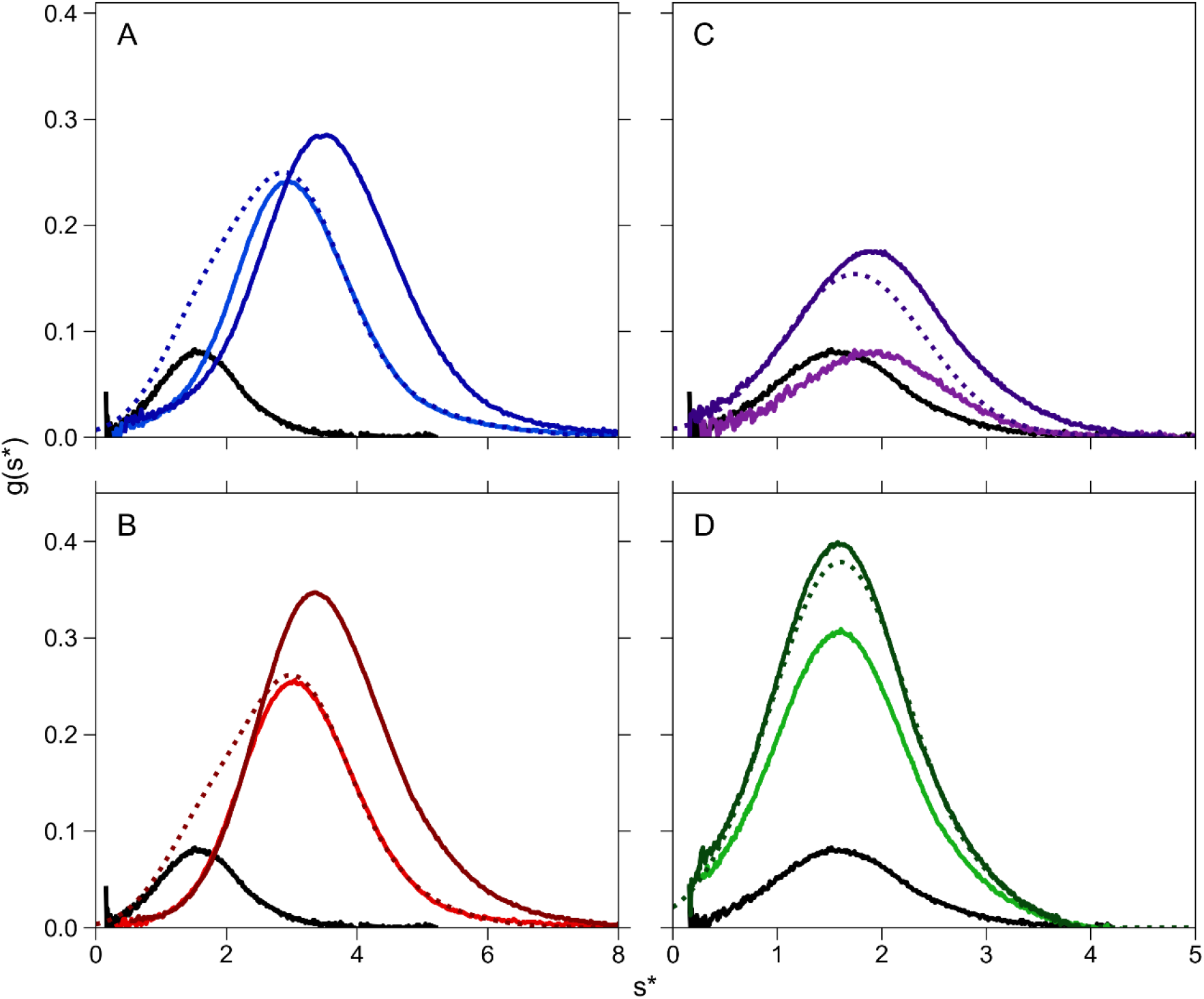
Shifted g(s*) distributions when titrating uOmpA_171_ with FkpA, sFkpA, and N-FkpA indicate binding. Distributions of 5 μM uOmpA_171_ (black) and 30 μM of FkpA construct were added to calculate the distribution for non-interacting species (dashed lines). Experimental reaction boundaries (solid lines) indicate that FkpA, sFkpA, and N-FkpA bind uOmpA_171_ at these concentrations. C-FkpA does not. A) fl-FkpA (blue), B) sFkpA (red), C) N-FkpA (purple), and D) C-FkpA (green).

## Discussion

OMP biogenesis requires a network of chaperones to keep uOMPs in an unfolded but folding-competent state in the aqueous environment of the periplasm. SurA, Skp, DegP, Spy, and FkpA have all been implicated in this periplasmic chaperone network,^8,9^ but the function of FkpA with respect to the OMP biogenesis pathway has been less investigated as compared to other chaperones (Figure 7). Here we show that FkpA increases the folded population of OmpA_171_ into diC_11_PC LUVs after one hour but decreases the observed rate of OmpA_171_ folding. To accomplish these seemingly contradictory functions, FkpA maintains the unfolded population of OmpA_171_ in a soluble state at early-to-intermediate time points and prevents the lipid-mediated aggregation that would otherwise occur. As a result, the folded population increases after an hour of folding even though the observed OmpA_171_ folding rate decreases in the presence of FkpA. To our knowledge, this is the first study to demonstrate a direct effect of FkpA on OMP folding.

**Figure 7.**
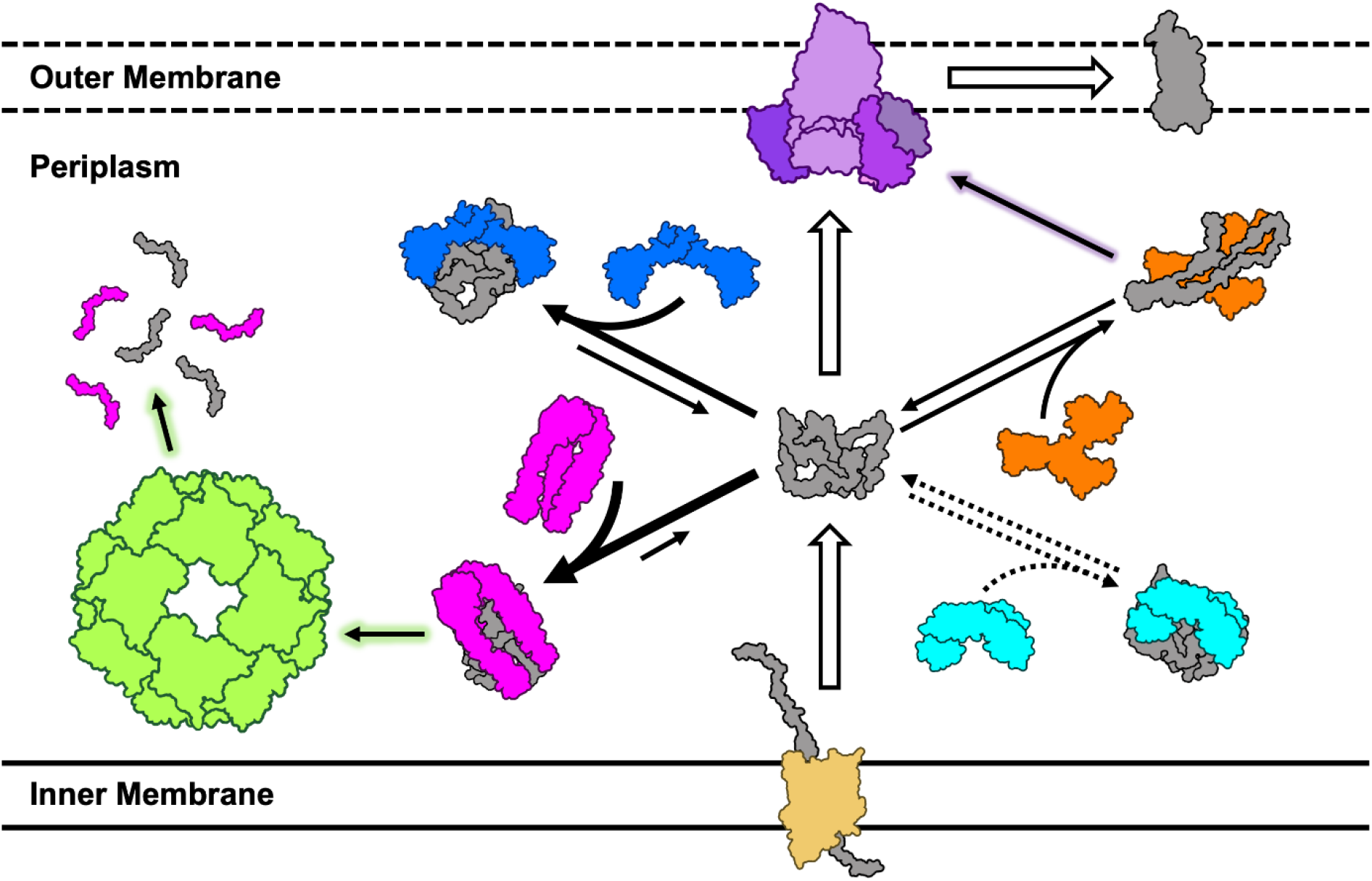
Schematic of the OMP biogenesis periplasmic chaperone network. As uOMPs are translocated into the periplasm, a network of periplasmic chaperones maintains uOMPs in an unfolded, but folding competent state before they fold into the membrane (white arrows) aided by the catalytic BAM complex (purple). FkpA (blue), Skp (pink), Spy (aqua), Deg (green), and SurA (orange) are members of this periplasmic network. Tight binders like Skp and FkpA favor the chaperone-uOMP complex to act as holdases that prevent uOMP aggregation. The tightly bound Skp complex can also be degraded by the proteases DegP (green arrow). Spy moonlights as an uOMP chaperone that binds some uOMPs with weak affinity (dotted arrows). SurA weakly binds uOMPs but contributes to the uOMP folding process by extending the unfolded polypeptides and handing off clients to the BAM complex (purple arrow). The thickness of the arrow indicates position of equilibrium in the binding reaction, not the flux of uOMPs toward a particular chaperone.

Despite the known redundancy encoded in the essentiality of the periplasmic chaperones, FkpA has a markedly different impact on OmpA_171_ folding as compared to the chaperones SurA or Skp. Further, FkpA does not behave as a folding catalyst like BAM. In contrast to FkpA, the addition of SurA to the folding reaction had no effect on OmpA_171_ folding, a finding that recapitulates previously published experiments investigating the folding of OmpA_171_ and PagP.^13,58^ This result in itself is somewhat surprising as SurA is thought to be the most important periplasmic chaperone in the OMP biogenesis pathway based on its phenotypic growth defect and the decrease in folded OMP levels upon SurA deletion.^48,49^ However, the function of SurA is thought to be linked to the BAM complex as evidenced by genetic and structural interactions between SurA and BAM.^48,59^ Additionally, recent literature supports a model where SurA binds extended conformations of uOMPs using localized and regulated binding regions, ^10,11,15,60,61^ leading to a weaker binding in the high nM to low μM range. ^13,54,55^ Quick client binding and release combined with a proposed interaction with and rate enhancement of BAM may explain why SurA can improve OMP folding in the presence of BAM but has little to no intrinsic ability to improve OMP folding efficiency by itself.^62,63^ Also in contrast to the functions revealed here for FkpA, the trimeric chaperone Skp keeps almost the entire population of OmpA_171_ in the unfolded state for the duration of the folding reaction, with only a small fraction of OmpA_171_ folding after a significant lag period. The effect of Skp is explained by its high affinity for uOMPs. Three monomers form a claw-like trimer with an internal cavity that sequesters a single uOMP chain, blocking aggregation with other chains.^64–66^ In this binding mechanism, a flexible Skp accommodates dynamic uOMPs through many non-specific interactions^17^ that contribute to an overall affinity in the low nM range.^18,55,56^ Because Skp binds uOMPs so tightly, they remain bound to the chaperone pool instead of folding into the lipid vesicles.

The fact that FkpA improves uOmpA_171_ folding efficiency over the course of an hour distinguishes FkpA from the other periplasmic chaperones, but FkpA is also not behaving as a folding catalyst like the BAM complex:^13,50,62^ FkpA does not enhance the rate of folding but instead diminishes it. Moreover, it is clear that the effect of FkpA on OmpA_171_ folding originates from its chaperone activity and not its PPIase activity. While FkpA is known to efficiently catalyze peptidyl-prolyl *cis/trans* isomerization in a range of peptide and full-length protein substrates,^24,26,31^ the sequence of OmpA_171_ does not include any *cis* prolines (SI Table S4). In fact, *cis* prolines are rare among both outer membrane and periplasmic proteins in general (SI Table S4), suggesting that the PPIase domain has been evolutionarily repurposed in periplasmic chaperones.

Attributing the effect of FkpA on uOmpA_171_ folding to a holdase activity is the simplest explanation. The strength of the interaction between FkpA and uOmpA_171_ may be “just right” to promote folding and prevent aggregation; a larger fraction of unfolded clients is kept soluble at intermediate time points and subsequently released and allowed to fold over the course of the reaction. Without increased molecular resolution, we cannot rule out the possibility that FkpA actively assists in uOMP folding perhaps by stabilizing secondary structures in the unfolded ensemble before membrane insertion. Further work to elucidate the mechanisms of FkpA client-binding and release and the structural features of an FkpA-uOMP complex are needed to fully explain the effect of FkpA on uOmpA_171_ folding.

In this work we also discovered that FkpA binds to uOMPs with an affinity that is intermediate between that of SurA and Skp. Therefore, in a direct competition of periplasmic chaperones, this implies that Skp ≥ FkpA > SurA in terms of individual binding affinity. One could argue that stable binding in the low nanomolar range is inconsistent with a chaperone function. However, as long as the rate of client release is fast enough, even small disassociation constants can be consistent with uOMP flux across the periplasm as has been previously shown.^6^ Even a disassociation rate on the order of seconds^22^ may be fast enough for facilitated diffusion of uOMPs without requiring additional factors to stimulate client release. However, we recognize that the periplasm is a complex environment that may also present competing reactions such as membrane charges, interactions with other periplasmic chaperones, the presence of the BAM complex, or self-regulation encoded in the conformational heterogeneity of FkpA. It is also possible that FkpA-uOMP complexes have the capacity to be fully degraded by DegP as has recently been shown for tightly-bound Skp-uOMP complexes (Figure 7).^67^

Previous work has suggested that the chaperone function of FkpA is localized to either the N-terminal dimerization^26,27^ or the C-terminal PPIase domain,^24,28^ but we find that neither domain alone exhibits chaperone function nor high affinity uOmpA_171_ binding. Some binding of uOmpA_171_ to N-FkpA was observed even though it was 10-fold weaker than the binding of the full-length protein, which may explain why N-FkpA prevented protein aggregation in some *in vitro* assays but was unable to rescue periplasmic stress when introduced into a *ΔfkpA ΔdegP* deletion strain.^27^ Results from the domain deletion constructs indicate that both domains in the full-length protein are required for full chaperone activity and tight client-binding. This is also consistent with the pAzF crosslinking assays, which indicate that the binding interface on FkpA spans the N- and C-domains with high efficiency crosslinking residues localized to regions on the inner surfaces of both. By bringing the two PPIase domains in close proximity via dimerization of the N-domain, FkpA increases the available binding surface and locally increases the number of interactions that contribute to the overall binding affinity. It should be noted that only the chaperone function of FkpA is dependent on both structural domains as the C-domain alone catalyzes peptidyl-prolyl isomerization reactions.^24,26^ Due to the extensive binding interface on the chaperone, the FkpA-uOmpA_171_ interaction is highly sensitive to the concentration of denaturant in solution. From the urea dependence of the binding constant, we estimate that approximately 22,000 Å^2^ of accessible surface area is buried when uOmpA_171_ binds FkpA (which is roughly equivalent to the ΔASA upon folding of α-chymotrypsin).^52^ Parameters like ΔASA and s_20,w_ of the complex obtained from the SV-AUC experiments are useful to constrain possible structures of an FkpA-uOMP complex.

The distinct effect of FkpA on OmpA_171_ folding requires the full-length, dimeric protein and suggests FkpA uses a strategy distinct from periplasmic chaperones SurA and Skp to directly impact uOMP folding. The intermediate-to-tight binding mediated by a large binding surface on the chaperone appears to a modified holdase strategy of aggregation mediation. In the context of the periplasmic network, the presence of FkpA ensures a robust response to stress, including environmental stresses encountered by virulent strains, by complementing the other unique roles of Skp and SurA (Figure 7, green and purple arrows). Continued work toward obtaining structural information on the FkpA_2_*OmpA_171_ complex and exploring the conformation heterogeneity of FkpA will help elucidate the full mechanism of FkpA chaperone function. As FkpA chaperones many unfolded, partially folded, or misfolded proteins other than its known OMP clients (OmpC and OmpA_171_), continued discovery of additional FkpA clients will help elucidate how a potent ATP-independent chaperone like FkpA balances the trade-off between specificity and promiscuity and functions to manage OMP biogenesis under periplasmic stress conditions. Our work indicates it functions as a modified holdase that is optimized for client release over a time frame that enhances folding efficiency. This modified holdase function of FkpA distinguishes it from other periplasmic chaperones but complements their functions for robust outer membrane biogenesis.

## Materials and Methods

### Expression and Purification of Proteins

Cytoplasmic expression and purification of soluble chaperones SurA, Skp, and all FkpA constructs is detailed in the supplemental information. Purification of OmpA_171_ from inclusion bodies can also be found in the supplemental information.

### Outer Membrane Protein Folding Assays

The preparation of diC_11_PC LUVs and execution of OmpA_171_ folding assays were modified from those conducted in Burgess *et al*.^68^ and Danoff *et al*.^40^ For a full description of these experiments, refer to the supplemental information.

### Sedimentation Velocity Analytical Ultracentrifugation Binding Titrations and Analysis

The oligomeric states of each FkpA construct and their binding affinities for uOmpA_171_ were determined using sedimentation velocity analytical ultracentrifugation (SV-AUC). Samples of each FkpA construct (FkpA, sFkpA, N-FkpA, and C-FkpA) were prepared at a series of concentrations between 1 and 150 μM in Phosphate Buffer (20 mM sodium phosphate, pH 8) supplemented with 1 M urea. All concentrations are the total monomer concentrations. A concentration series of FkpA in Phosphate Buffer was also prepared to verify that 1 M urea has no effect on the oligomerization of FkpA. To determine binding affinities, 5 μM uOmpA_171_ was titrated with each FkpA construct from 1 nM to 150 μM of the chaperone. FkpA titrations were repeated in 0.8 M, 0.9 M and 1.2 M urea. The sedimentation of 5 μM uOmpA_171_ has been previously shown to be monomeric and monodisperse in 1 M urea, ^5,51^ which was verified by spinning 5 μM OmpA_171_ in Phosphate Buffer plus 1 M urea in triplicate.

All SV-AUC experiments were performed using a Beckman XL-A ultracentrifuge (Beckman Coulter) and cells with 1.2 mm double-sector epoxy centerpieces and sapphire windows. Each sample was spun at 20 °C using a 4-hole, An-Ti60 rotor and speed of 50000 rpm for up to 200 scans. Radial scans were acquired with 0.003 cm radial steps in continuous mode with zero time interval between scans. A wavelength of 230, 250, or 280 nm was chosen for each experiment such that the sample absorbance was always in the linear optical range of 0.1 – 1.0 absorbance units. Extinction coefficients (SI Table S3) at 230 nm and 250 nm were obtained from the slope of calibration curves created with a series of known protein concentrations (data not shown). Prior to starting each run, the rotor was temperature equilibrated in the instrument for at least 60 minutes.

All SV-AUC data was analyzed using dc/dt+^69^ and SEDANAL.^70^ Sedimentation coefficient distributions (g(s*) distributions) were created using dc/dt+ and were corrected using the appropriate densities (ρ), viscosities (η), and partial specific volumes 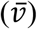 for each buffer and protein construct calculated using SEDNTERP (SI Table S4).^71^ Full FkpA concentration series were first globally fit in SEDANAL to determine the oligomeric state of each construct alone. The best models and statistics describing these fits can be found in Table 1. To obtain the best fit, a small (< 1%) fraction of irreversible FkpA aggregate was included in the model when fitting FkpA and sFkpA concentration series.

Binding titrations were globally fit in SEDANAL to determine disassociation constants (K_d_) and the sedimentation coefficients of the chaperone-uOmpA_171_ complex. FkpA and sFkpA titrations were fit to a uOmpA_171_ + FkpA_2_ ↔ uOmpA_171_*FkpA_2_ binding model including the irreversible FkpA aggregate at < 1% of the total protein concentration. The N-FkpA titration could also be fit to the simple uOmpA_171_ + N-FkpA_2_ ↔ uOmpA_171_*N-FkpA_2_ model, but the C-FkpA titration was best described by two ideal species that do not interact and thus no binding could be observed. The sedimentation coefficients of uOmpA_171_and each FkpA construct was held fixed at its previously determined value when fitting the corresponding binding titration. For all global fits, goodness-of-fit was assessed by minimization of residuals reflected in the root-mean-square deviation (RMSD), by the randomness of the residuals, and narrowness of the 95% confidence interval on fitted parameters. Confidence intervals were determined using the F-statistic module in SEDANAL (Table 2). Sedimentation coefficients from SEDANAL fits were corrected to 20 °C in water using Equation 1.

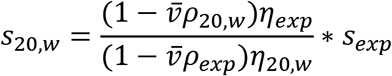

### Chaperone Competition Pulldown Assays

To explore how FkpA binds uOmpA_171_ in the context of other periplasmic chaperones, we designed a chaperone competition experiment based on a previously published assay.^21^ Specifically, we investigated whether FkpA can outcompete SurA or Skp for uOMP binding. To compare SurA and FkpA, 20 μM His-tagged SurA was mixed with 5 μM uOmpA_171_ in Phosphate Buffer containing 1 M urea. A total volume of 1 mL was loaded onto 500 μL of Ni-NTA Sepharose High Performance resin in a 1.5 mL Eppendorf tube and allowed to equilibrate for 5 minutes. The flowthrough was collected by centrifugation for 5 min at 14000 rpm in an Eppendorf tabletop centrifuge. All subsequent wash and elution steps were also equilibrated on the resin for 5 min and collected by centrifugation. Then 1 mL of 40 μM untagged FkpA in Phosphate Buffer containing 1 M urea was added to the resin to compete with SurA in binding uOmpA_171_ (FkpA Wash). A second wash using 1 mL of Buffer A followed the FkpA Wash. Protein remaining bound to the column was eluted by adding 2 x 1 mL of Buffer B to the column. To compare FkpA and Skp, 30 μM His-tagged Skp was mixed with 5 μM uOmpA_171_ in Phosphate Buffer containing 200 mM NaCl and 1 M urea. The pulldown assay was performed using the above protocol except that Skp bound to the column was eluted by adding 4 x 1 mL of Buffer B to the column followed by a single elution with 1 mL of Buffer C.

SDS-PAGE samples were prepared by combining 15 uL of each flowthrough, wash, or elution with 5 μL of 4 X SDS Loading Buffer and heating at 95 °C for 5 minutes. Samples were run on 4 - 20% gradient precast gels (Mini-PROTEAN TGX; Bio-Rad) run at a constant voltage of 200 V for 35 min at room temperature. The intensity of the uOmpA_171_ band in each lane was determined using densitometry analysis in ImageJ. The intensities in all wash steps and all elution steps were added to the cumulative values reported in Figure 3DE.

### Photo-crosslinking Assays

Samples of 50 μM of each FkpA pAzF variant alone and 50 μM of the FkpA pAzF variant mixed with 5 μM uOmpA_171_ in Phosphate Buffer plus 1 M urea were prepared in a total volume of 40 μL then split into two aliquots of 15 μL each. One of the two aliquots (UV-treated sample) was then irradiated with UV light (wavelength, λ = 254 nm) for 5 min using a Spectroline MiniMax UV Lamp (11-992-662; Fisher) to cross-link the species in solution. Each control (no UV exposure) and UV-treated aliquot was denatured by addition of 5 μL of 4 X SDS Loading Buffer and heating at 95 °C for 5 minutes. Samples were subjected to SDS-PAGE using a 4 to 20% gradient precast gel (Mini-PROTEAN TGX; Bio-Rad) run at a constant voltage of 200 V for 35 min at room temperature. Using ImageJ, densitometry analysis on the uOmpA_171_ bands in the control (I_-UV_) and UV-treated (I_+UV_) conditions was utilized to quantitate crosslinking efficiency (XL Efficiency) according to Equation 2. Crosslinking efficiency values were corrected for the amount of uOmpA_171_ lost (~11%) when mixed with wild-type, full-length FkpA (not containing pAzF) and subject to UV irradiation (correction factor = CF). A representative SDS-PAGE gel for photo-crosslinking between each FkpA pAzF variant and uOmpA_171_ is shown in SI Figure S9.

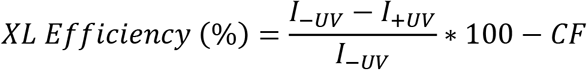

All other materials and methods can be found in the supplemental information.

## Supplemental Materials Description

Supplementary materials are included as a single document Supplement.pdf. It includes four tables and ten figures in additional to supplementary methods and results.

## Acknowledgements

We thank the Center for Molecular Biophysics for providing facilities and resources. The authors thank lab members for helpful discussions. This work was funded by National Science Foundation (NSF) grants MCB 1931211 (K.G.F). T. D., D. C. M., and A. M. P. were supported by National Institutes of Health (NIH) training grant T32-GM008403. A. M. P. was supported by NSF Grant DGE 1232825.

## Abbreviations and Symbols

ASA: accessible surface area
AUC: analytical ultracentrifugation
BAM: β-barrel assembly machine
CD: circular dichroism
FkpA: FK506-binding protein A
IDR: intrinsically disordered region
LUV: large unilamellar vesicles
Ni NTA: Nickel nitrilotriactetic acid
OMP: outer membrane protein
pAzF: *para*-azido phenylalanine
PPIase: peptidyl-prolyl isomerase
RMSD: root mean square deviation
uOMP: unfolded outer membrane protein
s_20,w_: sedimentation coefficient corrected to water at 20 °C
SDS-PAGE: sodium dodecyl sulfate – polyacrylamide gel electrophoresis
σ^E^: sigma E stress response
Skp: seventeen kilodalton protein
Spy: spheroplast protein Y
SurA: survival factor A
SV: sedimentation velocity
UV: ultraviolet
XL: crosslinking

