## Supplemental Information for "FkpA Enhances Membrane Protein Folding using an Extensive Interaction Surface"

### Supplement Table of Contents

#### Supplemental Methods

*Purification of $OmpA_{171}$ from Inclusion Bodies*

Cloning of $OmpA_{171}$ into a pET11a vector with ampicillin resistance is described in Danoff, 2011.[1] Glycerol stocks of $OmpA_{171}$ plasmid transformed into HMS174(DE3) cells that had been stored at -80 C were used to start 5 mL terrific broth (TB, Fisher) overnight cultures containing 100 ug/mL ampicillin (Sigma). The following day, overnight cultures were used to inoculate 500 mL of TB containing 100 ug/mL ampicillin, and cultures were grown at 37 °C with shaking until reaching an optical density at 600 nm ($OD_{600}$) of 1.0. Protein expression was induced by supplementing cultures with 1mM isopropyl-β-D1-thiogalactopyranoside (IPTG) (ThermoScientific) and incubating them at 37 °C with shaking for an additional 4-6 hours. Cells were then harvested by centrifugation in a Beckman J2-MI centrifuge using a JA-10 rotor at 5000 rpm and 4 °C for 30 min. Cell pellets were stored at -20 C until lysis.

A pellet from a 500 mL growth was resuspended in 25 mL of OMP Lysis Buffer (50 mM Tris, 40 mM ethylenediaminetetraacetic acid (EDTA), pH 8) and subsequently lysed using an Avestin Emulsiflex homogenizer. After lysis, Brij-L23 (Sigma) was added to a final concentration of 0.1%, and full lysates were centrifuged (Beckman J2-MI centrifuge, JA-10 rotor) at 5000 rpm and 4 °C for 30 minutes. Pellets containing the isolated inclusion bodies were washed twice by resuspending in 25 mL of OMP Wash Buffer (10 mM Tris, 1 mM EDTA, pH 8) and subsequent centrifugation under the same conditions. The purified inclusion bodies were resuspended a final time in 10 mL of OMP Wash Buffer and were aliquoted into 1 mL portions in 1.5 mL Eppendorf tubes before centrifuging a final time at 14,000 rpm and room temperature for 30 minutes using a table-top centrifuge (Eppendorf). The supernatant was discarded, and inclusion body pellets were stored at -20 °C.

An inclusion body aliquot was resuspended in 1 mL of the appropriate buffer (OMP Folding Buffer – 10 mM borate, 2 mM EDTA, pH 10 used in folding assays or Phosphate Buffer – 20 mM sodium phosphate, pH 8 used in sedimentation velocity experiments) then pelleted at 14,000 rpm and room temperature for 5 minutes using a table-top centrifuge (Eppendorf). The inclusion body pellet was dissolved in 1.2 mL of 8 M Urea in the appropriate buffer background (OMP Folding Buffer or Phosphate Buffer) by incubation at room temperature for >15 minutes. Once the pellet was fully dissolved, contaminating nucleic acids were removed by centrifugation at 14,000 rpm and room temperature for 5 minutes using a table-top centrifuge (Eppendorf). The supernatant was carefully pipetted off, and final sample purity and concentration were assessed by collecting a UV-visible absorbance wavelength spectrum. Samples were deemed pure of nucleic acid contaminants if the A260/A280 ≤ 0.6, and protein concentration was determined using the theoretical extinction coefficient of 45,090 $M^{-1}$ $cm^{-1}$ calculated using the Edelhoch method[2] in SEDNTERP[3] (SI Table S3). Protein was diluted with 8 M Urea in the appropriate buffer (OMP Folding Buffer or Phosphate Buffer) to a final concentration of approximately 80 μM and then aliquoted and stored at -80 °C until use.

*Expression and Purification of all SurA and FkpA Constructs*

The *E. coli* FkpA gene was inserted into the pET28b vector containing kanamycin resistance between the Nde I and Xho I restriction sites and was followed by a C-terminal Tobacco Etch Virus (TEV) protease cleavage site and a 6-Histidine tag. The SurA plasmid was designed by inserting the *E. coli* gene for SurA with a C-terminal 6-Histidine tag into the pET28b vector between restriction sites Nde I and BamHI. Both FkpA and SurA constructs lack a periplasmic signal sequence in order to overexpress in the cytoplasm, but residue numbering in this paper includes the signal sequences. FkpA deletion constructs (sFkpA, N-FkpA, C-FkpA) were cloned using the In-Fusion HD Cloning Plus CE method (Takara). Primers are listed in Table S2. Stellar cells were transformed with the PCR product by heat shock and plated grown overnight on Luria Broth (LB) plates with 50 μg/ml kanamycin at 37 °C. Plasmid DNA was extracted from single colonies with the GeneJET Plasmid Miniprep Kit and sequenced to validate the construct sequences. Plasmids were transformed into *E. coli* HMS174(DE3) cells via electroporation for protein expression and stored at −80 °C as glycerol stocks.

Glycerol stocks were used to inoculate 5 mL TB growths containing 50 mg/mL kanamycin. These cultures were grown overnight at 37 ˚C with shaking and then diluted into 500 mL TB also supplemented with 50 ug/mL kanamycin. Cultures were grown at 37 °C with shaking until reaching an $OD_{600}$ of 0.6-0.8 when overexpression was induced by the addition of 1mM IPTG (ThermoScientific). Cells containing SurA and all FkpA construct plasmids were expressed overnight (>16 hours) at 37 °C. The following morning cell pellets were collected by centrifugation at 5000 rpm and 4 °C for 30 min in a Beckman J2-MI centrifuge with a JA-10 rotor. Cell pellets were frozen and stored at -20 °C until lysis.

All FkpA constructs and SurA were purified according to the following protocol. Previously thawed cell pellets were solubilized in 25 mL of Buffer A (20 mM sodium phosphate, 500 mM NaCl, 20 mM imidazole, pH 8) supplemented with an EDTA-free protease inhibitor tablet (Pierce). Resuspended cells were mechanically lysed using an Emulsiflex homogenizer (Avestin) then centrifuged at 17000 rpm and 4 °C for 30 minutes in a Beckman J2-MI centrifuge using a JA-21 rotor. The supernatant was filtered using a 0.22 μm syringe-top filter (Millipore) before loading onto a Ni-NTA Sepharose High Performance bench-top column (GE Healthcare) pre-equilibrated in Buffer A. Protein bound to the column was first washed and denatured with 40 mL of Buffer A plus 6 M urea, then washed and refolded with an additional 20 mL of Buffer

A. Proteins were eluted in 5 mL fractions using 25 mL of Buffer B (20 mM sodium phosphate, 500 mL NaCl, 300 mM imidazole, pH 8), and the column was finally washed with 25 mL of Buffer C (20 mM sodium phosphate, 500 mL NaCl, 500 mM imidazole, pH 8). Fractions containing protein were pooled and concentrated to approximately 1 mL using 15 mL Amicon spin concentrators (Millipore) with a 30 kDa molecular weight cut-off (MWCO) for FkpA, sFkpA, and SurA or 10 kDa MWCO for N-FkpA and C-FkpA. As a final purification and buffer exchange step, proteins were run over a Superdex 75 Increase 10/300 GL column (GE Healthcare) in Phosphate Buffer (20 mM sodium phosphate, pH 8) using a fast performance liquid chromatography (FLPC) system (BioRad) at room temperature. The purity of eluted fractions was assessed by sodium dodecyl sulfate polyacrylamide gel electrophoresis (SDS-PAGE), and pure fractions were pooled. Final protein concentrations were determined using the absorbance at 280 nm and the theoretical extinction coefficients calculated using the Pace method[4] in SEDNTERP.[3] All extinction coefficients are listed in Table S3. Proteins were aliquoted and stored at -80 °C until use.

To remove the C-terminal His-tag from FkpA for the chaperone competition assays, tobacco etch virus (TEV) protease was added to purified His-tagged FkpA in a 20:1 molar ratio of FkpA:TEV protease. The mixture was dialyzed at room temperature in Buffer A supplemented with 3 mM dithiothreitol (DTT) for 48 hours before switching into a DTT-free Buffer A for an additional 24 hours. Cleaved protein was collected as the flowthrough from a Ni-NTA Sepharose High Performance bench-top column and was buffer exchanged into Phosphate buffer using a Superdex 75 Increase 10/300 GL column (GE Healthcare) on the FPLC system.

*Incorporation of* para-*azido phenylalanine (pAzF)*

A library of 24 FkpA *para*-azido phenylalanine (pAzF) variants was created by incorporating Amber stop codons (TAG) at individual positions in the gene using Infusion Cloning (Takara) All primers are listed in Table S2. Surface exposed sites were chosen using the 1Q6U crystal structure. For pAzF incorporation, HMS cells were transformed with both the FkpA pAzF plasmid and the pEVOL-pAzF plasmid, which encodes the amber-suppressor tRNATyr(CUA) from *Methanocaldococcus jannaschii* and its cognate aminoacyl-tRNA synthetase under an arabinose inducible promoter.[5] pEVOL-pAzF was a gift from Peter Schultz (Addgene plasmid # 31186 ; http://n2t.net/addgene:31186 ; RRID:Addgene_31186). Bacteria growth and protein expression followed the same protocol as used for wild-type, full-length FkpA except that cultures were supplemented with 1 mM pAzF and 0.2 % arabinose in addition to IPTG at the induction step. All subsequent lysis, purification, and storage procedures were identical to those used for wild-type, full-length FkpA.

*Expression and Purification of Skp*

The nomenclature NZ100 defines the pOPINE construct harboring Skp with an 8-histidine C-terminal tag that was cloned as described previously.[6] BL21 (DE3) pLysS cells in a glycerol stock containing NZ100 were used to inoculate 5 mL of sterile LB containing 60 μg/mL ampicillin, 34 μg/mL chloramphenicol, and 1% glucose. Overnight cultures grown at 37° C were diluted into 500 mL of 2XYT media in baffled flasks and grown in the presence of antibiotic and glucose to an $OD_{600}$ of 0.6-0.8. Upon reaching the appropriate OD, cells were induced with 1 mM IPTG and grown for 5 hours at 37 °C. Cells were pelleted by centrifugation at 5000 rpm and 4 °C for 30 min in a Beckman J2-MI centrifuge with a JA-10 rotor, and cell pellets were frozen and stored at -20 °C until lysis. Thawed cell pellets were solubilized in 25 mL of Buffer A supplemented with an EDTA-free protease inhibitor tablet (Pierce). Resuspended cells were mechanically lysed using an Emulsiflex homogenizer (Avestin) then centrifuged at 17000 rpm and 4 °C for 30 minutes in a Beckman J2-MI centrifuge using a JA-21 rotor. The supernatant was filtered using a 0.22 μm syringe-top filter (Millipore) before loading onto a Ni-NTA Sepharose High Performance bench-top column (GE Healthcare) pre-equilibrated in Buffer A. Protein bound to the column was first washed and denatured with 40 mL of Buffer A plus 6 M urea, then washed and refolded with an additional 20 mL of Buffer A. Proteins were eluted in 5 mL fractions using 25 mL of Buffer B, and the column was finally washed with 25 mL of Buffer C. Fractions containing protein were pooled and concentrated to approximately 1 mL using 15 mL Amicon spin concentrators (Millipore) with a 30 kDa MWCO. As a final purification and buffer exchange step, proteins were run over a Superdex 75 Increase 10/300 GL column (GE Healthcare) in 20 mM sodium phosphate, 20 mM NaCl, pH 8 using a FLPC system (BioRad) at room temperature. The purity of eluted fractions was assessed by sodium dodecyl sulfate polyacrylamide gel electrophoresis (SDS-PAGE), and pure fractions were pooled. Final protein concentrations were determined using the absorbance at 280 nm and the theoretical extinction coefficients calculated using the Pace method[4] in SEDNTERP.[3] All extinction coefficients are listed in Table S3. Skp was stored at 4 °C for up to two weeks after the final size exclusion purification step.

*Preparation of Large Unilamellar Vesicles*

Phosphostidylcholine lipids (Avanti Polar Lipids) dissolved in chloroform were dried to a thin film in glass vials (12.5 mg/vial) using a gentle flow of nitrogen. The vials were further evacuated overnight to remove any residual solvent and stored at -20 C until use. To prepare the large unilamellar vesicles (LUVs), the lipid films were first reconstituted in 500 uL of OMP

Folding Buffer (10 mM borate, 2 mM EDTA, pH 10) to a final concentration of 25 mg/mL and left to incubate, with occasional vortexing, at room temperature for at least 30 minutes. The reconstituted lipids were then extruded 15 times through a 0.1 μm filter using a mini-extruder (Avanti Polar Lipids) to form the LUVs. All folding assays in this study were performed using 1,2-diundecanoyl-$_{sn}$-glycero-3-phosphocholine (diC$_{11}$PC) LUVs. LUVs created by this process have been previously shown to form stable unilamellar vesicles using transmission electron microscopy.[7]

*OmpA$_{171}$ Folding Assays*

Folding of OmpA$_{171}$ into diC$_{11}$PC LUVs both in the presence and absence of chaperones was monitored using SDS-PAGE. In a 3 mL glass cuvette, OMP Folding Buffer (10 mM borate, 2 mM EDTA, pH 10) containing 1 M urea, diC$_{11}$PC LUVs, and chaperone, if present, were combined in a final volume of 500 μL and left to incubate at 37 °C in a custom-built stirring incubator (Aviv Medical) for 10 minutes. Folding assays were performed using a total lipid concentration of 3.2 mM. All FkpA constructs were added to a final monomer concentration of 40 μM, SurA was added to a final concentration of 20 μM, and Skp was added to a final monomer concentration of 30 μM. To initiate the folding assay, OmpA$_{171}$ was rapidly diluted from an 80 μM stock in 8 M urea, 10 mM borate, 2mM EDTA, pH 10 to a final concentration of 4 μM, which equates to an 800:1 lipid:protein ratio. Small aliquots of the folding reaction were removed and quenched with 4X SDS gel loading buffer to a final concentration of 1X to prevent further folding at timepoints 5, 12, 25, 40, 60, 120, 180, 300, 600, 1200, 2400, and 3600 seconds after folding initiation. Quenched aliquots at each timepoint remained at room temperature until 10 μL was loaded onto a precast gel (Mini-PROTEAN TGX, Bio-Rad). Additional aliquots were also removed and quenched at 1 min and 1 hour after folding initiation. These additional aliquots were heated at 95 °C for 5 min to fully unfold all $OmpA_{171}$ prior to gel-loading. All samples were analyzed by SDS−PAGE the same day they were collected. Folding reactions containing $OmpA_{171}$, $OmpA_{171}$ + FkpA, $OmpA_{171}$ + sFkpA, $OmpA_{171}$ + N-FkpA, $OmpA_{171}$ + SurA, and OmpA + Skp were assayed using 12% acrylamide gels. Reactions monitoring $OmpA_{171}$ folding in the presence of C-FkpA were assayed using 10% acrylamide gels. All gels were stained with Coomassie Blue R-350 (GE Healthcare) and scanned digitally. Densitometry was performed using ImageJ.[8]

Intensities of the folded and unfolded bands obtained from densitometric analyses were used to quantify the folded and unfolded populations of $OmpA_{171}$. Fraction folded and unfolded were calculated by dividing the intensity of the folded (F) and unfolded (U) bands respectively by the average intensity of the two boiled samples (B).

$$Fraction\ Folded = \frac{F}{B}$$

$$Fraction\ Unfolded = \frac{U}{B}$$

Because the summation of these two fractions do not add to unity, a third "elusive" fraction that does not migrate with either the folded or unfolded populations was accounted for using the following equation:

$$Fraction\ Elusive = \frac{(B - (F + U))}{B}$$

For all folding conditions, fractional populations are shown as the average of at least 7 independent replicates (n=7 for $OmpA_{171}$, $OmpA_{171}$+ SurA, $OmpA_{171}$+ Skp, and $OmpA_{171}$ + N-FkpA, n=8 for $OmpA_{171}$ + FkpA, n=9 for $OmpA_{171}$ + C-FkpA, n=12 for $OmpA_{171}$ + sFkpA). Error bars are ±1 standard deviation. Plots of fraction folded as a function of time were fit to a single exponential function of the form,

$$y = y_0 + Ae^{-kt}$$

where y is the fraction folded at a given time, $y_0$ is the fraction folded as time approaches infinity, k is the observed first-order rate constant, and A is the negative amplitude. Observed first-order rate constants ($k_{obs}$) are reported as the average of at least 7 independent replicates and error bars are ±1 standard deviation.

*Calculation of Theoretical Sedimentation Coefficients*

Ensembles of FkpA were creating using coarse-grained molecular dynamics simulations in CafeMol.[71] CafeMol models each amino acid as a bead at the $C_\alpha$ position and utilizes a Go-like potential in addition to excluded volume, hydrophobic, and electrostatic potentials during the simulation. Starting from the PDB file for the structure of FkpA (PDB 1Q6U),[27] N- and C-terminal tails in additional to an N-terminal methionine and C-terminal TEV protease site and 6X Histidine tag were added to the file in PyMol to match the sequence of expressed protein used in this study. Coarse-grained MD simulations were run using the Go-like, excluded volume, hydrophobic, and electrostatic potentials on the folded domains of FkpA, but the Go-like potential was removed from the intrinsically disordered regions at the N- and C-termini and from a region of helix III (113-117) that has previously been shown to exhibit increased dynamics by NMR.[28,30] Coarse-grained simulations were run on MARCC for 25000000 frames at a temperature of 310 K with a timestep of 0.2 and a hydrophobic parameter of 0.58. Every other frame of the last 2000 frames were taken to represent the ensemble. Side chains were reintroduced using PULCHRA,[72] and the final 1000 structures were relaxed by NAMD.[73] The $s_{20,w}$ of each frame was calculated using HullRad[74] to create a distribution of $s_{20,w}$ values, and the average $s_{20,w}$ was compared to experimental values (SI Figure S2, Table 2). Simulations to create structural ensembles were done on both monomeric and dimeric FkpA, sFkpA, and N-FkpA. Only monomeric C-FkpA was simulated.

*Circular Dichroism Spectroscopy*

Circular dichroism (CD) wavelength scans from were collected of each FkpA deletion construct. Each protein was diluted to 1-3 μM in 20 mM sodium phosphate, pH 8 or 20 mM sodium phosphate, pH 8, 1 M urea, and wavelength scans were collected from 200-300 nm at 25 C. Chemical denaturation of 2.7 μM FkpA in Phosphate buffer was performed by titrating in 8 M urea in Phosphate buffer using a MicroLab 500 series automatic titrator (Hamilton). To check for path independence, 2.7 μM of FkpA in 8 M urea in Phosphate buffer was titrated with Phosphate buffer. Unfolding and folding of FkpA was monitored by the respective decrease or increase in signal at 222 nm. All CD experiments were done using 1 cm quartz cuvettes and a Model 410 Aviv CD spectrometer.

*OMP Proline Composition Analysis*

The most abundant OMPs and periplasmic proteins have been previously identified[9,10] and are summarized in Table S4. We enumerated the number of prolines in each of these proteins via a Python script. We then utilized the dihedral measurement tool in PyMol to measure the backbone omega (ω) dihedral angle for all proline residues identified in our analysis. Four atoms define the ω dihedral angle: the carbonyl O from residue$^{Pro-1}$, the backbone C from residue$^{Pro-1}$, the N from the proline, and the Ca from the proline. Proline residues with an w angle of ± 180° ± 15° are classified as *cis* (SI Table S1).

#### Supplemental Results

*Oligomeric States of FkpA Constructs*

It has been established that FkpA is dimeric above 1 μM[11,12] but to verify the oligomeric state of FkpA under our experimental conditions (20 mM sodium phosphate, pH 8, 20 °C, in 0 M Urea and 1 M Urea) at we collected a concentration series of sedimentation velocity analytical ultracentrifugation experiments (SV-AUC) and globally fit the data from the full range of concentrations (1-100 μM) to a model in SEDANAL (SI Figure S3).[13] The FkpA concentration series in both 0 M and 1 M urea was best described by a single, dimeric species containing a small fraction of irreversible aggregate (< 1%). A sedimentation coefficient of 2.77 (2.76-2.78, 95% confidence interval) was obtained from the global fit to data collected in 1 M urea (Table 1). After correcting to 20 °C in water, an $s_{20,w}$ = 3.05 Svedbergs matched the $\langle s_{20,w} \rangle$ = 2.83 ±0.15 Svedbergs calculated using HullRad[14] on an ensemble of dimeric structures from coarse-grained molecular dynamics simulations (Table 1, SI Figure S2).[15]

Like FkpA, a series of SV-AUC data at sFkpA concentrations between 1-100 μM, in 1 M urea and at 20 °C is best described by the sFkpA dimer (Table 1, SI Figure S2). C-FkpA is monomeric under these same experimental conditions (Table 1, SI Figure S2), which agrees with previously published work.[12] However, while it has been previously reported that N-FkpA exists in a monomer-dimer equilibrium with a $K_{dim}$=10-100 μM,[16] the N-FkpA concentration series was best fit by a single ideal species, and the $s_{20,w}$ = 2.01 (2.00-2.02) obtained matches the values calculated for dimeric N-FkpA (Table 1, SI Figure S2).

**Supplemental Figures**

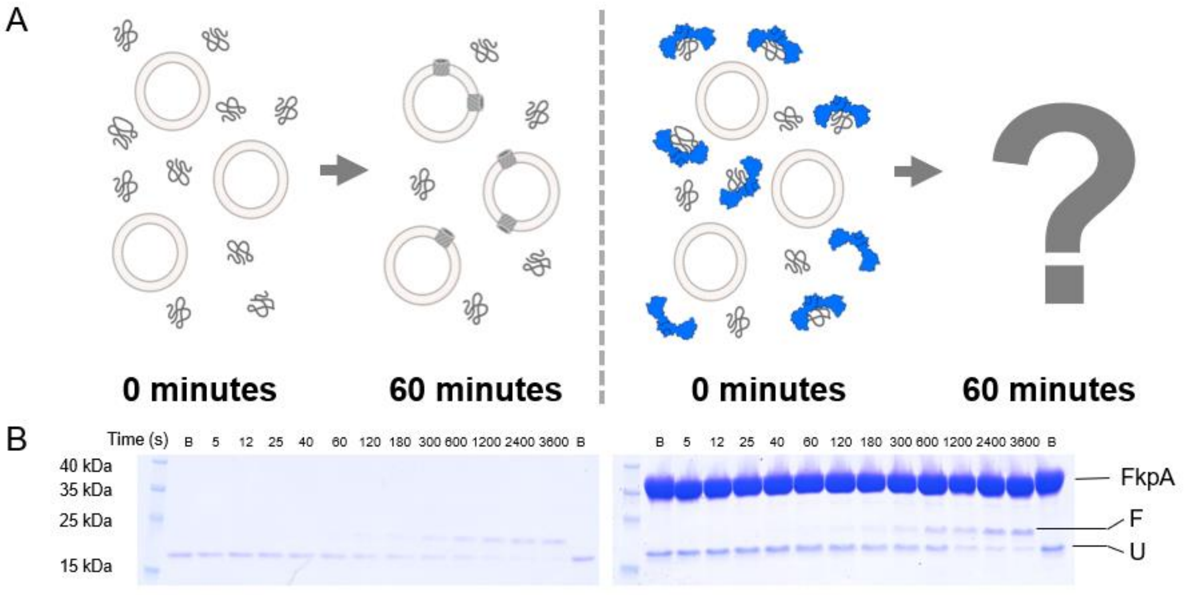

*Figure S1. OmpA$_{171}$ Folding Assays in the Absence and Presence of FkpA.* A) Schematic of uOmpA$_{171}$ folding into LUVS in the absence of chaperone (left). The effect of FkpA on folding was probed by adding chaperone prior to folding initiation (right). B) Representative SDS-PAGE images of OmpA$_{171}$ folding assays in the absence of chaperone (left) and in the presence of FkpA (right). The folded (F) and unfolded (U) populations appear as two distinct bands when samples are denatured by SDS but not heated and can be quantified by densitometry in ImageJ. The unfolded population of OmpA$_{171}$ migrates more quickly than the folded population.

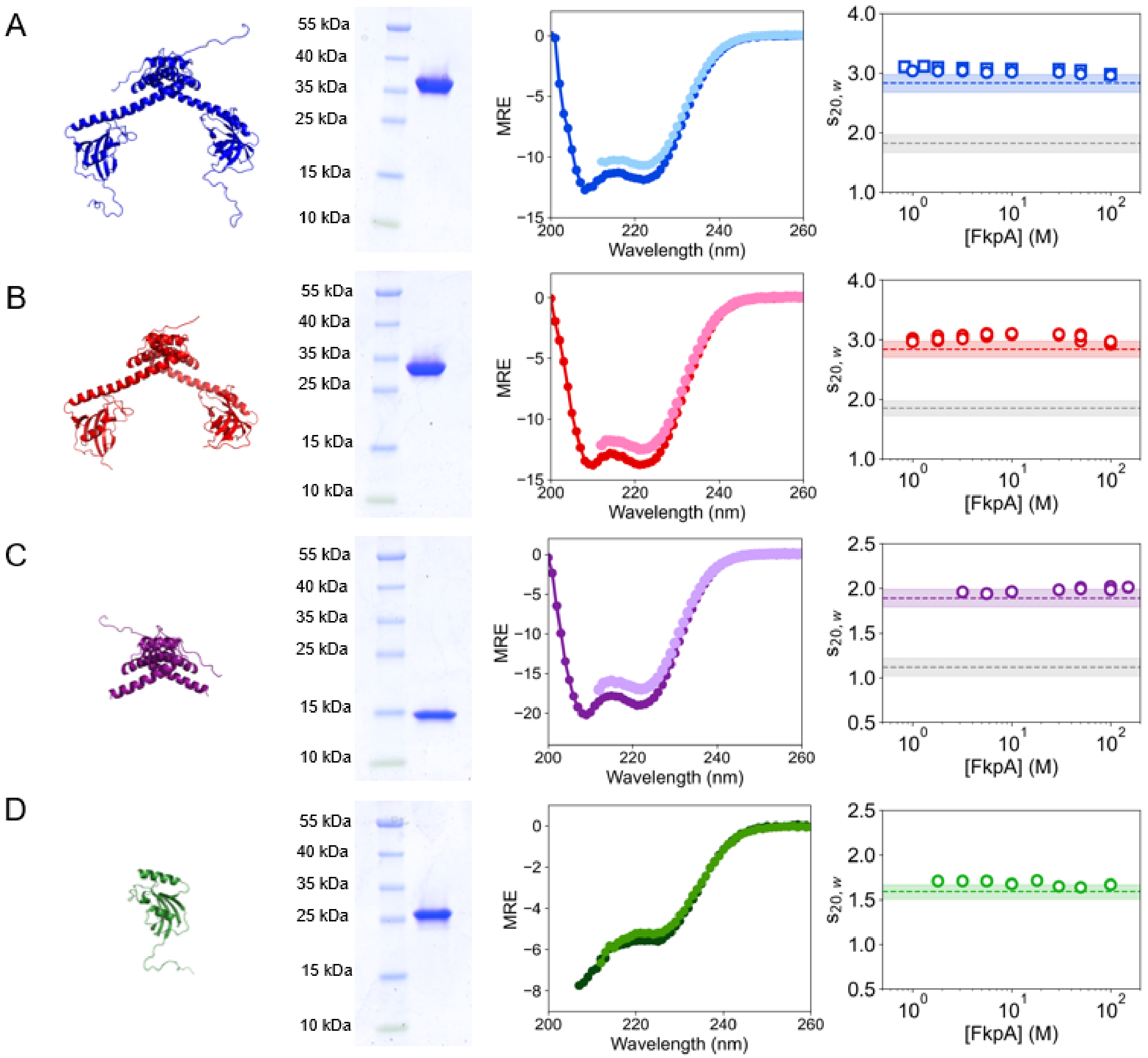

*Figure S2. Purity, Foldedness, and Oligomeric State of FkpA Constructs.* A) FkpA, B) sFkpA, C) N-FkpA, and D) C-FkpA. (Left) Ribbon diagrams of each FkpA construct were created in PyMol starting from PDB 1Q6U. (Left Center) SDS-PAGE confirms that each construct is pure and (Right Center) CD wavelength spectra show that each construct is folded at 0 M (dark color) and 1 M (light color) urea. (Right) Experimental $s_{20,w}$ values at a range of monomer protein concentrations compared to the theoretical values of different oligomers. Measurements made in 0 M urea are shown as open squares. Measurements in 1 M urea are shown as open circles. Sedimentation coefficients calculated using HullRad on an ensemble of structures are shown as dotted lines while the filled regions represent the standard deviation in the calculated value. For FkpA, sFkpA, and N-FkpA, gray lines represent the theoretical sedimentation coefficient of the monomer, and colored lines represent the theoretical sedimentation coefficient of the dimer. Based on these results, FkpA, sFkpA, and N-FkpA are dimers across the experimental concentration range. C-FkpA is monomeric.

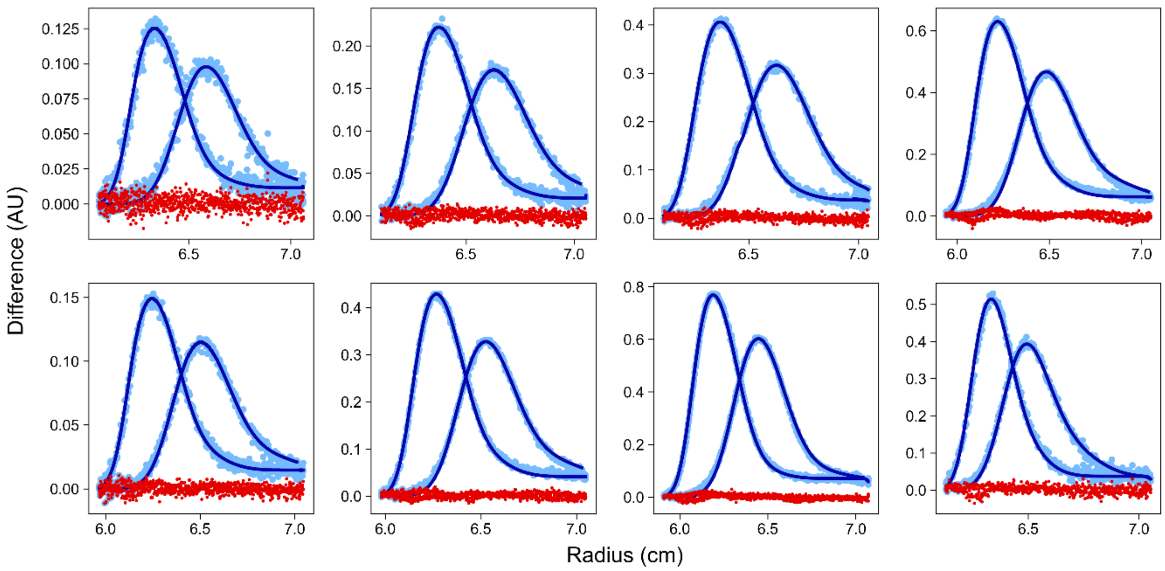

*Figure S3. SEDANAL fit of the FkpA concentration series to a single ideal dimer model*.

Concentrations of FkpA increase from 1 μM (top left) to 100 μM (bottom right) in Phosphate Buffer containing 1 M urea. Experimental data is depicted as light blue points, the fit is a dark blue line, and residuals are in red. The fit $s_{20,w}$ = 3.05 (3.04-3.06), and the model fits with an RMSD = 0.0067.

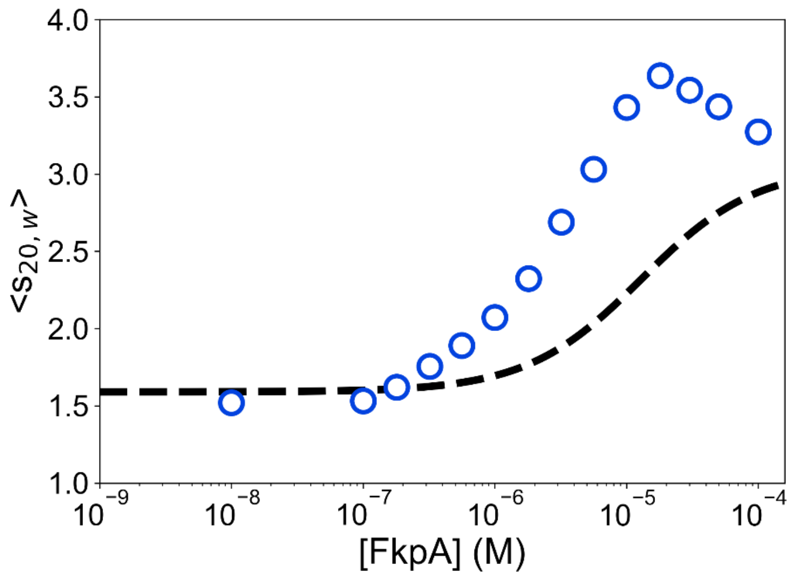

*Figure S4. Clear uOmpA$_{171}$ binding to FkpA is demonstrated by a shifted <s$_{20,w}$>.* uOmpA$_{171}$ (5 μM) was titrated with FkpA from 0.01-100 uM in 1 M urea, 20 mM sodium phosphate, pH 8. The average $s_{20,w}$ (<s$_{20,w}$>) of each mixture was calculated in dcdt+. If uOmpA$_{171}$ and FkpA did not interact, the reaction boundary and <s$_{20,w}$> would change according to the concentration-weighted average of the two species (see dashed black line). Instead, experimental values (blue open circles) shift to larger <s$_{20,w}$>, indicating binding. After binding saturates, the <s$_{20,w}$> decreases toward the sedimentation coefficient of dimeric FkpA (approx. 3 Svedbergs) due to the presence of free, excess FkpA. Here FkpA concentrations are monomeric concentrations.

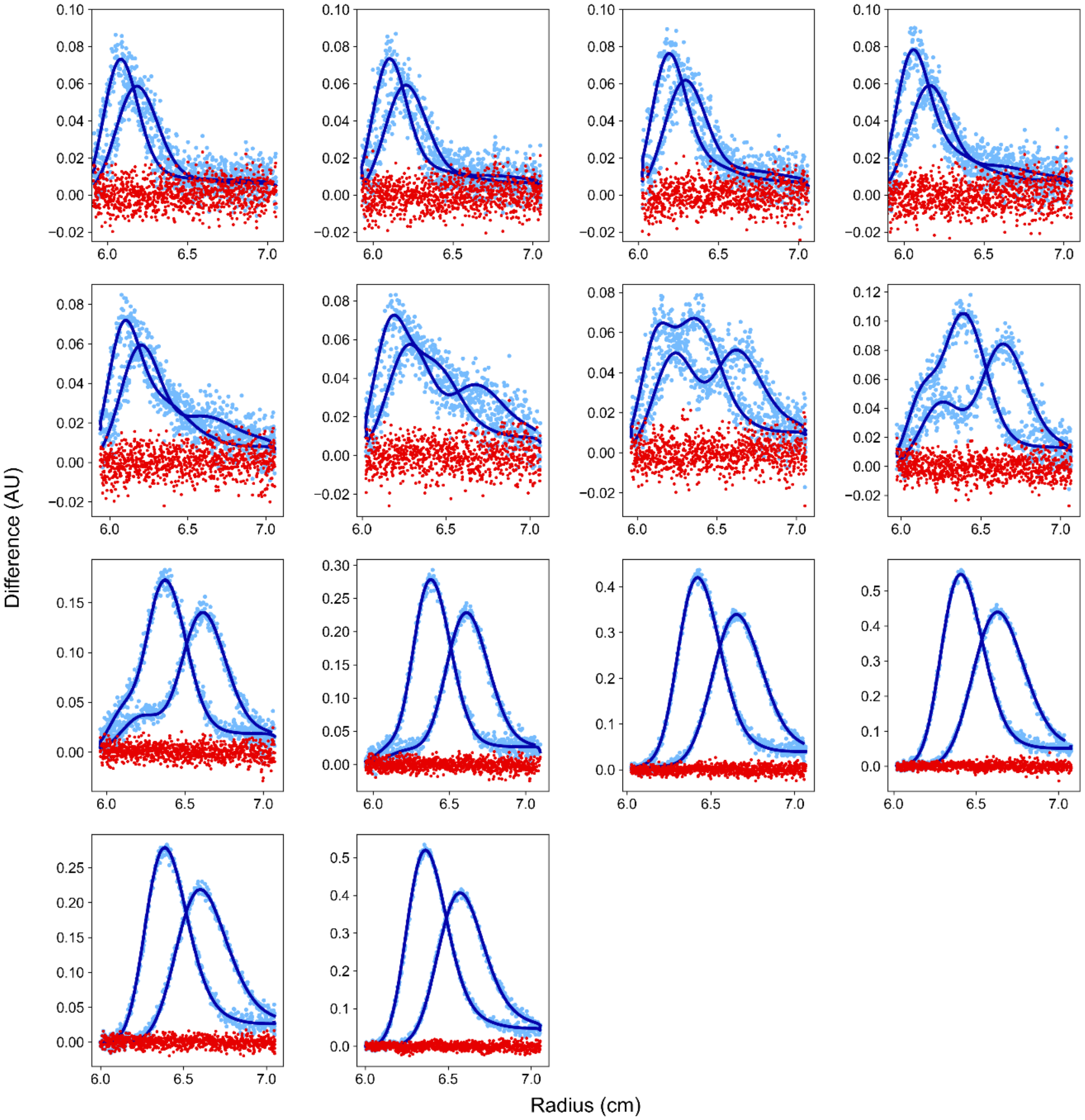

*Figure S5. SEDANAL fit of FkpA titrating $uOmpA_{171}$ using the model $FkpA_2$ + $uOmpA_{171}$ ↔ $FkpA_2$*$uOmpA_{171}$*. Concentrations of FkpA increase from 10 nM (top left) to 100 µM (bottom right) while the $uOmpA_{171}$ concentration was held constant at 5 µM in Phosphate Buffer containing 1 M urea. Experimental data is depicted as light blue points, the fit is a dark blue line, and residuals are in red. The complex $s_{20,w}$ = 3.8 (3.68-3.92) Svedbergs and $K_d$ = 1.1 (0.6-1.8) µM. The model fits with an RMSD = 0.0071.

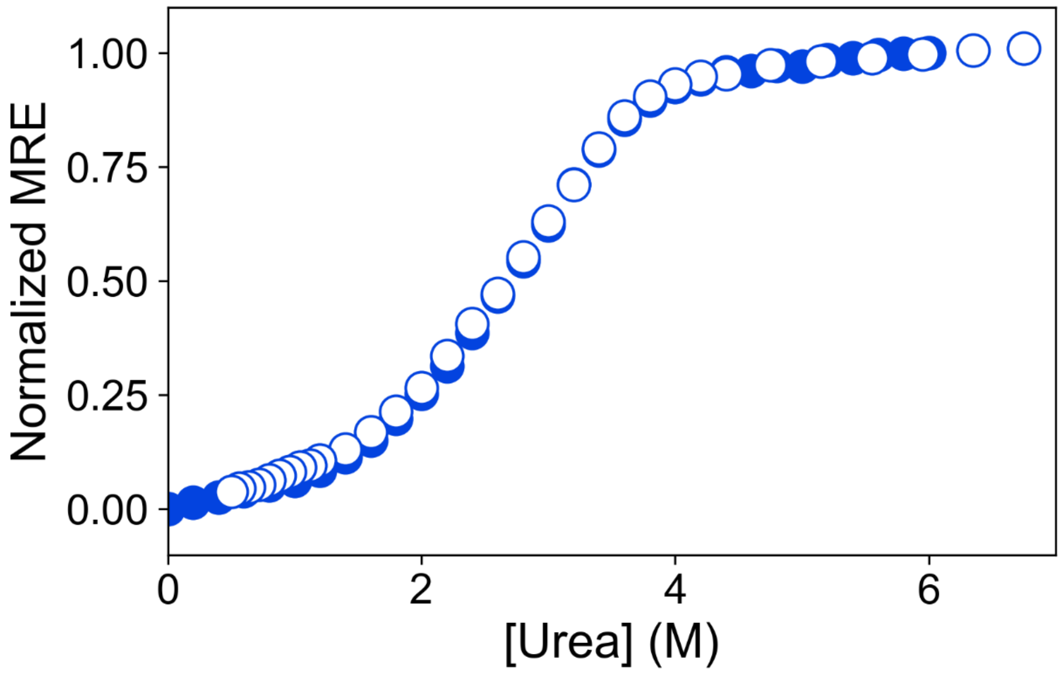

*Figure S6. Chemical denaturation of FkpA.* Loss of secondary structure was followed by the change in signal at 222 nm as 2.7 μM FkpA was titrated with 8 M urea. Folding of FkpA in Phosphate Buffer at 25 °C is reversible (unfolding curve in filled circles, refolding curve in open circles). The unfolded fraction of FkpA becomes significant above 1.5 M urea, precluding any binding titrations at higher urea concentrations.

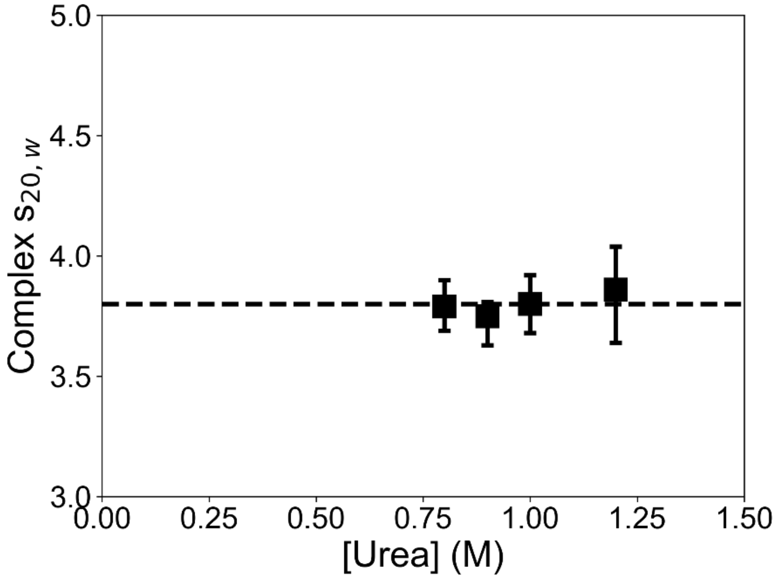

*Figure S7. Complex $s_{20,w}$ is mostly insensitive to the urea concentration.* The sedimentation coefficient of the FkpA*uOmpA$_{171}$ complex varies little between 0.8-1.2 M urea. The complex $s_{20,w}$ at 1.0 M urea is shown as a horizontal dashed line for reference.

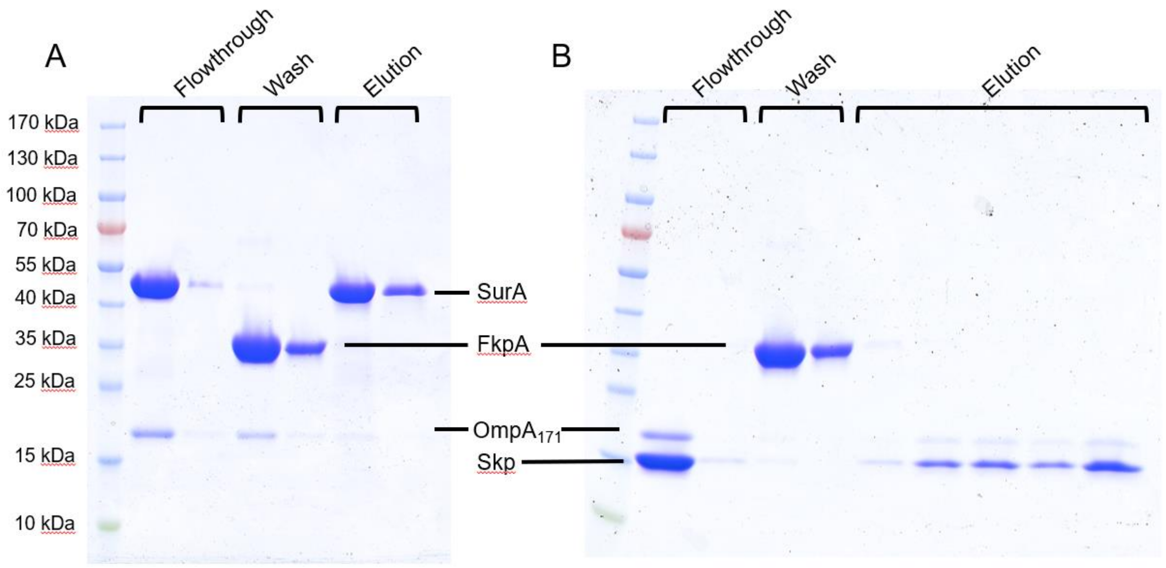

*Figure S8. FkpA can outcompete SurA but not Skp for uOmpA$_{171}$ binding*. A) SDS-PAGE gel of all flowthrough, wash, and elution fractions after pre-equilibrating the Ni resin with SurA and uOmpA$_{171}$ and washing with untagged FkpA. B) SDS-PAGE gel of all flowthrough, wash, and elution fractions after pre-equilibrating the Ni resin with Skp and uOmpA$_{171}$ and washing with untagged FkpA.

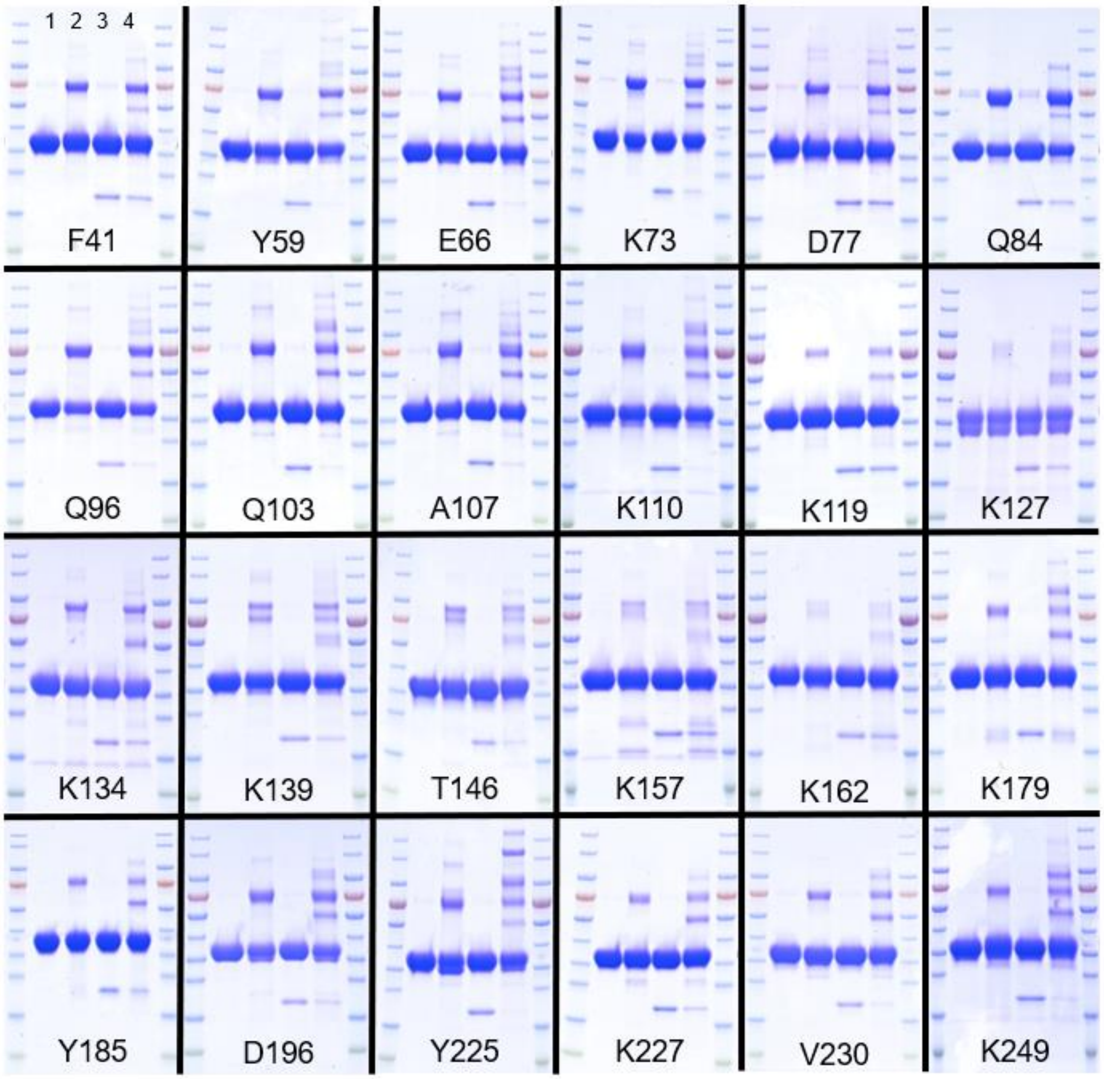

*Figure S9. Representative SDS-PAGE images of all FkpA pAzF crosslinking variants*. The noncanonical amino acid *para*-azido phenylalanine (pAzF) was incorporated into FkpA at 24 surface exposed positions. The position replaced by pAzF is written on each gel image. Each variant was crosslinked to uOmpA$_{171}$ by UV light, and crosslinking was quantified using SDS-PAGE. Lane 1 – 50 µM FkpA pAzF variant, no UV exposure. Lane 2 – 50 µM FkpA pAzFvariant with 5 minutes UV exposure. Lane 3 – 50 µM FkpA pAzF and 5 µM uOmpA$_{171}$, no UV exposure. Lane 4 – 50 µM FkpA pAzF variant and 5 µM uOmpA$_{171}$ with 5 minutes UV exposure. Crosslinking efficiency was quantified as the loss of band intensity for uOmpA$_{171}$ after UV exposure. Crosslinked complexes appear as higher molecular weight complexes.

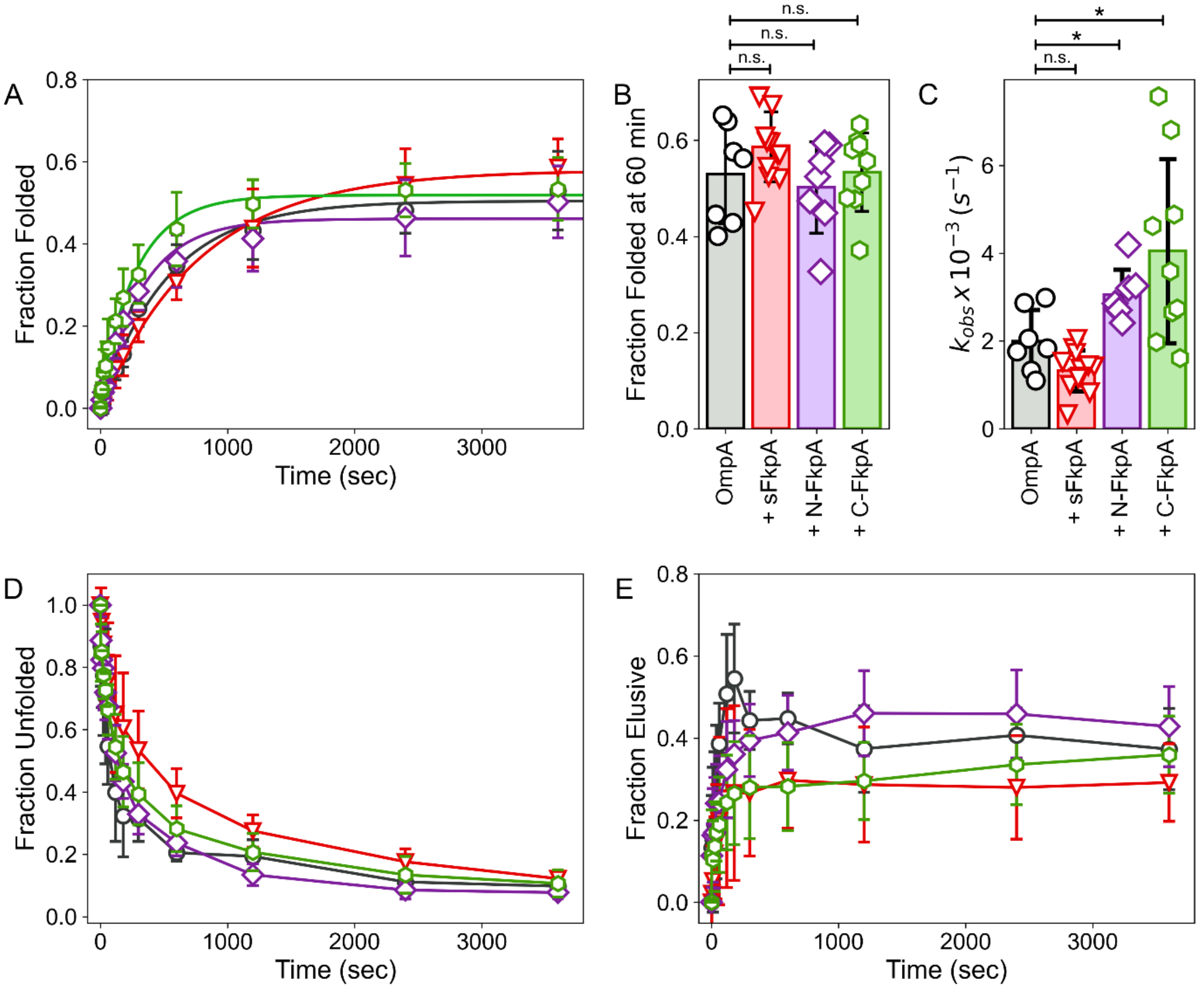

*Figure S10. Domain deletion constructs do not exhibit the same chaperone function as full-length protein*. A) Fraction of $OmpA_{171}$ folded into $diPC_{11}$ LUVS as a function of time (0-3600 s) in the presence of sFkpA (red triangles, n=12), N-FkpA (purple diamonds, n=7), C-FkpA (green hexagons, n=9) or without any chaperone (black cicles, n=7). Error bars represent ±1 standard deviation. Lines are single exponential fits to the average data. B) Comparison of the fraction of $OmpA_{171}$ folded at 60 min. C) Comparison of the observed rate constants when individual datasets are fit to single exponential functions. (* $p < 0.05$, independent t-test with unequal variances) D) Unfolded fraction of $OmpA_{171}$ as a function of time. E) Elusive fraction as a function of time. Error bars represent ±1 standard deviation.

#### Supplemental Tables

*Table S1. Proline contents of the most abundant OMPs and periplasmic proteins*

| | | PDB | Total # Residues | Total # Prolines | # *cis* Prolines |
|---|---|---|---|---|---|
| Most Abundant OMPs | LamB | 1MAL | 421 | 8 | 1 |
| | $OmpA_{171}$ | 1QJP | 171 | 8 | 0 |
| | OmpC | 2J1N | 346 | 3 | 0 |
| | OmpF | 3K19 | 340 | 4 | 0 |
| | OmpT | I78 | 297 | 9 | 0 |
| | OmpW | 2F1V | 191 | 6 | 0 |
| | OmpX | 1QJ9 | 148 | 4 | 1 |
| Most Abundant Periplasmic Proteins | AgP | 1NT4 | 391 | 24 | 0 |
| | GlnH | 1WDN | 226 | 7 | 1 |
| | HdeA | 1DJ8 | 89 | 4 | 0 |
| | HdeB | 2XUV | 79 | 5 | 0 |
| | MalE | 1ANF | 370 | 21 | 0 |
| | OppA | 3TCH | 517 | 32 | 1 |
| | RbsB | 1URP | 271 | 9 | 0 |

*Table S2. All primers used in this study.* Any stop codons introduced into the sequence are in bold.

| FkpA Construct | Residues Removed/Modified | Primers |
|---|---|---|

| | | |
|---|---|---|
| sFkpA | Δ26-34 | Forward: CCCATATGGCTGACAGCAAAGCAGCGT<br><br>Reverse: TGTCAGCCATATGGGTATATCTCCTTCTTAAA |
| | Δ253-270 | Forward: CAGCGCCGCTCGAGAGCAGCGGTGAGG<br><br>Reverse: TCTCGAGCGGCGCTGGTTTCACATCC |
| N-FkpA | Δ122-270 | Forward: AAGACGCGCTCGAGAGCAGCGGTGAGG<br><br>Reverse: TCTCGAGCGCGTCTTTTTCCATCTTCGCC |
| C-FkpA | Δ26-121 | Forward: CAAAACCTGCTGATAACGAAGCAAAAGGTA<br><br>Reverse: TATCAGCAGGTTTTGCAGCTTCAGCC |
| F41pAzF | F41 | Forward: AGCAGCG**TAG**AAAAATGACGATCAGAAATCAGC<br><br>Reverse: TTTTT**CTA**CGCTGCTTTGCTGTCAGC |
| Y59pAzF | Y59 | Forward: GGTCGT**TAG**ATGGAAAACTCTCTAAAAGAACAAG<br><br>Reverse: TTCCAT**CTA**ACGACCCAGCGAGGCAC |
| E66pAzF | E66 | Forward: TCTAAAA**TAG**CAAGAAAAACTGGGCATCAAACTG<br><br>Reverse: TCTTG**CTA**TTTTAGAGAGTTTTCCATGTAACGA |
| K73pAzF | K73 | Forward: GGGCATC**TAG**CTGGATAAAGATCAGCTGATCGCTG<br><br>Reverse: TCCAG**CTA**GATGCCCAGTTTTTCTTGTTCTTT |
| D77pAzF | D77 | Forward: GGATAAA**TAG**CAGCTGATCGCTGGTGTTCA<br><br>Reverse: AGCTG**CTA**TTTATCCAGTTTGATGCCCAGT |
| Q84pAzF | Q84 | Forward: TGGTGTT**TAG**GATGCATTTGCTGATAAGAGCAAAC<br><br>Reverse: GCATC**CTA**AACACCAGCGATCAGCTGATC |
| Q96pAzF | Q96 | Forward: CTCCGAC**TAG**GAGATCGAACAGACTCTACAAGCA<br><br>Reverse: ATCTC**CTA**GTCGGAGAGTTTGCTCTTATCAGCA |
| Q103pAzF | Q103 | Forward: GACTCTA**TAG**GCATTCGAAGCTCGCGTG |

| | | |
|---|---|---|
| | | Reverse: AATGC**CTA**TAGAGTCTGTTCGATCTCTTGGTCG |
| A107pAzF | A107 | Forward: ATTCGAA**TAG**CGCGTGAAGTCTTCTGCTCAG<br>Reverse: ACGCG**CTA**TTCGAATGCTTGTAGAGTCTGTTCG |
| K110pAzF | K110 | Forward: TCGCGTG**TAG**TCTTCTGCTCAGGCGAAGATGG<br>Reverse: GAAGA**CTA**CACGCGAGCTTCGAATGC |
| K119pAzF | K119 | Forward: GATGGAA**TAG**GACGCGGCTGATAACGAAGC<br>Reverse: GCGTC**CTA**TTCCATCTTCGCCTGAGCAG |
| K127pAzF | K127 | Forward: CGAAGCA**TAG**GGTAAAGAGTACCGCGAGAAATTTG<br>Reverse: TTACC**CTA**TGCTTCGTTATCAGCCGCG |
| K134pAzF | K134 | Forward: CCGCGAG**TAG**TTTGCCAAAGAGAAAGGTGTG<br>Reverse: GCAAA**CTA**CTCGCGGTACTCTTTACCTTTT |
| K139pAzF | K139 | Forward: CAAAGAG**TAG**GGTGTGAAAACCTCTTCAACTGG<br>Reverse: ACACC**CTA**CTCTTTGGCAAATTTCTCGCGG |
| T146pAzF | T146 | Forward: CTCTTCA**TAG**GGTCTGGTTTATCAGGTAGTAGAAG<br>Reverse: AGACC**CTA**TGAAGAGGTTTTCACACCTTTCTC |
| K157pAzF | K157 | Forward: AGCCGGT**TAG**GGCGAAGCACCGAAAGAC<br>Reverse: TCGCC**CTA**ACCGGCTTCTACTACCTGATAAACC |
| K162pAzF | K162 | Forward: AGCACCG**TAG**GACAGCGATACTGTTGTAGTGAAC<br>Reverse: CTGTC**CTA**CGGTGCTTCGCCTTTACCG |
| K179pAzF | K179 | Forward: CGACGGT**TAG**GAGTTCGACAACTCTTACACCCGT<br>Reverse: AACTC**CTA**ACCGTCGATCAGCGTACCTTTG |
| Y185pAzF | Y185 | Forward: CAACTCT**TAG**ACCCGTGGTGAACCGCTT<br>Reverse: CGGGT**CTA**AGAGTTGTCGAACTCTTTACCGTCG |

| | | |
|---|---|---|
| D196pAzF | D196 | Forward: CCGTCTG**TAG**GGTGTTATCCCGGGTTGG<br>Reverse: ACACC**CTA**CAGACGGAAAGAAAGCGG |
| Y225pAzF | Y225 | Forward: ACTGGCT**TAG**GGCAAAGCGGGTGTTCCG<br>Reverse: TTGCC**CTA**AGCCAGTTCTGGTGGAATAACC |
| K227pAzF | K227 | Forward: TTACGGC**TAG**GCGGGTGTTCCGGGGATCC<br>Reverse: CCCGC**CTA**GCCGTAAGCCAGTTCTGGTGG |
| V230pAzF | V230 | Forward: AGCGGGT**TAG**CCGGGGATCCCACCGAATTC<br>Reverse: CCCGG**CTA**ACCCGCTTTGCCGTAAGC |
| K249pAzF | K249 | Forward: GGATGTG**TAG**CCAGCGCCGAAGGCTGAT<br>Reverse: GCTGG**CTA**CACATCCAGCAGCTCTACGTCA |

Residue numbering in this paper includes the 25 aa periplasmic signal sequences even though these are not in the cytoplasmically expressed proteins

*Table S3. Properties of all proteins used in study*

| | | | | Extinction coefficient (ε) ($M^{-1}$ $cm^{-1}$) | | |
|---|---|---|---|---|---|---|
| Protein | # Residues | MW (Da) | $\bar{\nu}$ (mL/g) | 280 nm | 230 nm | 250 nm |
| $OmpA_{171}$ | 172 | 18875 | 0.7219 | 45090 | --- | 14400 |
| FkpA | 267 | 27856 | 0.7341 | 17420 | 140000 | 7120 |
| sFkpA | 240 | 26157 | 0.7344 | 17420 | 143000 | 6060 |
| N-FkpA | 118 | 12908 | 0.7217 | 4470 | 62300 | --- |
| C-FkpA | 149 | 18984 | 0.7388 | 14440 | 105000 | 4860 |
| SurA | 417 | 46274 | --- | 29160 | --- | --- |

| | | | | | | |
|---|---|---|---|---|---|---|
| Skp | 149 | 16792 | --- | 1490 | --- | --- |

*All values were calculated for monomeric proteins with sequence files that include the N-terminal Methionine and any Histidine tags

*Table S4. Properties of all buffers used in SV-AUC experiments.* Densities and viscosities were calculated in SEDNTERP.

| Temperature (°C) | Buffer | [Urea] (M) | ρ (g/mL) | η (cP) |
|---|---|---|---|---|
| 20 | 20 mM sodium phosphate, pH 8 | 0 | 1.00108 | 1.0116 |
| | | 0.8 | 1.01331 | 1.0410 |
| | | 0.9 | 1.01488 | 1.0406 |
| | | 1.0 | 1.01645 | 1.0487 |
| | | 1.2 | 1.01960 | 1.0593 |
| | Water | --- | 0.99823 | 1.0020 |

**Supplemental Information References**

1. Danoff, E. J. & Fleming, K. G. The soluble, periplasmic domain of OmpA folds as an independent unit and displays chaperone activity by reducing the self-association propensity of the unfolded OmpA transmembrane β-barrel. *Biophys. Chem.* **159**, 194–204 (2011).
2. Edelhoch, H. Spectroscopic Determination of Tryptophan and Tyrosine in Proteins. *Biochemistry* **6**, 1948–1954 (1967).
3. Laue, T. M., Shah, B. D., Ridgeway, T. M. & Pelletier, S. L. Computer-aided interpretation of analytical sedimentation data for proteins. in *Analytical Ultracentrifugation in Biochemistry and Polymer Science* (eds. Harding, S., Rowe, A. & Hoarton, J.) 90–125 (Royal Society of Chemistry, 1992).
4. Pace, C. N., Vajdos, F., Fee, L., Grimsley, G. & Gray, T. How to measure and predict the molar absorption coefficient of a protein. *Protein Sci.* **4**, 2411–2423 (1995).
5. Chin, J. W. *et al.* Addition of p-azido-L-phenylalanine to the genetic code of Escherichia coli. *J. Am. Chem. Soc.* **124**, 9026–9027 (2002).
6. Moon, C. P., Zaccai, N. R., Fleming, P. J., Gessmann, D. & Fleming, K. G. Membrane protein thermodynamic stability may serve as the energy sink for sorting in the periplasm. *Proc. Natl. Acad. Sci. U. S. A.* **110**, 4285–4290 (2013).
7. Danoff, E. J. & Fleming, K. G. Novel kinetic intermediates populated along the folding pathway of the transmembrane β-barrel OmpA. *Biochemistry* **56**, 47–60 (2017).
8. Schneider, C. A., Rasband, W. S. & Eliceiri, K. W. NIHImage to ImageJ: 25 years of image analysis. *Nat. Methods* **9**, 671–676 (2012).
9. Han, M.-J., Kim, J. Y. & Kim, J. a. Comparison of the large-scale periplasmic proteomes

of the Escherichia coli K-12 and B strains. *J. Biosci. Bioeng.* **117**, 437–42 (2014).
